# TransPepMS: A Tool to Verify Peptides Absent from Reference Proteomes

**DOI:** 10.64898/2026.10.20.760571

**Authors:** S. Yasin Tabatabaei Dakhili, Olivier Hinse, Leonard J. Foster, Dieter Brömme

## Abstract

Cells generate peptides absent from reference proteomes, including products of reverse proteolysis and gene fusions. These are an increasingly pursued class of targets in autoimmune disease, cancer vaccines and T-cell therapies. The sequence is itself the discovery, yet a convincing spectral match does not establish it, because an ordinary genome-encoded peptide of the same mass can explain the spectrum equally well. Independent benchmarking has identified this ambiguity as a dominant source of error in reported non-canonical identifications. Here, we introduce TransPepMS, which adjudicates each proposed assignment and returns a verdict of retained, unresolved, or rejected. For example, on adjudicating 1,154 reported proteasome-spliced immunopeptides, only 53 survived same-mass competition, whereas 91 of 99 insulin- spike hybrids were retained and all 46 planted canonical peptides were rejected, showing that TransPepMS discriminates genuine non-canonical peptides rather than merely deflating claims. TransPepMS provides peptide validation before they guide biological interpretation and immune-target selection.

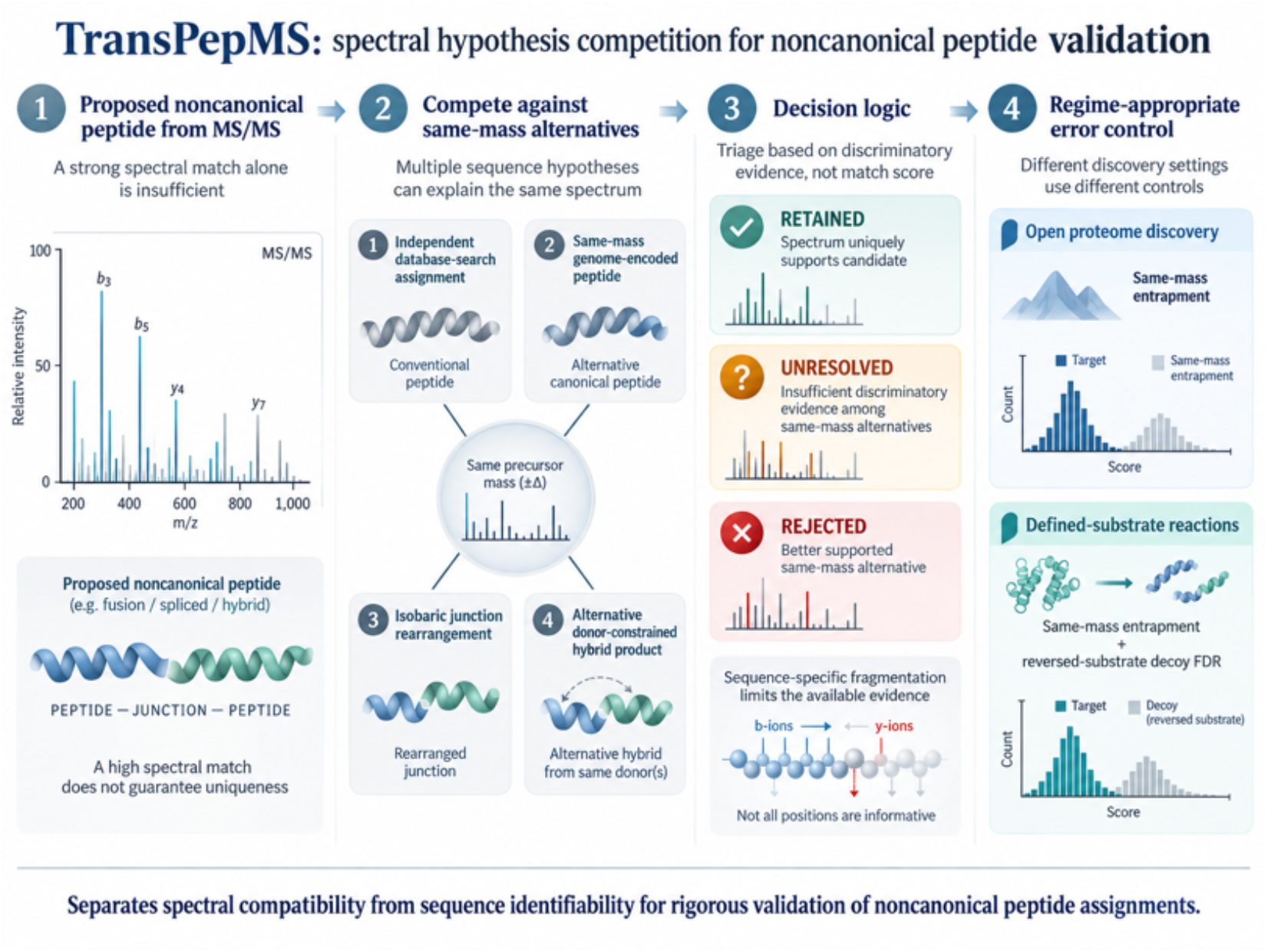

## Introduction

The functional and structural diversity of the proteome extends far beyond the encoding of the canonical genome through mechanisms such as ribosomal frameshifting and reverse proteolysis ^1–4^. Non-canonical peptides, those absent from reference proteomes, are of particular interest because they expand the repertoire of immune targets in cancer and autoimmunity. The biological importance of such sequences is exemplified by proteasome-generated spliced peptides, hybrid insulin peptides in Type 1 Diabetes (T1D), and various fusion-junction neoantigens ^5–16^. Uncovering these sequences links a candidate peptide directly to hypotheses regarding its biogenesis and immune recognition. However, these biological interpretations hinge entirely on correct sequence assignment: an incorrect assignment risks directing downstream research toward a non-existent sequence.

A convincing spectral match alone does not prove that a proposed non-canonical sequence is better supported than a plausible precursor-mass isobar ^2^. A standard genome-encoded peptide may explain the observed fragment ions equally well or better. Indeed, independent reanalyses of proteasome-splicing datasets exposed this fundamental ambiguity by reassigning spectra originally attributed to spliced peptides back to linear genome-encoded sequences ^17–19^. Beyond individual datasets, a community benchmark of reported non-canonical identifications found wide disagreement between studies and limited high-quality spectral support, underscoring the need for standardized, per-spectrum validation ^1^. The central challenge is therefore not whether a proposed sequence *can* match a spectrum, but whether the spectrum reliably *distinguishes* that sequence from all plausible alternatives. Without this distinction, assignment uncertainty propagates into flawed conclusions about the composition and biological origin of the peptide repertoire.

Existing analytical controls fail to resolve this ambiguity. Database searches evaluate only the candidate sequences contained within their specified search space ^20–22^; even custom databases expanded to include predicted fusion or spliced peptides ^23–25^ cannot evaluate alternatives they do not contain. Target-decoy approaches estimate overall false discovery rates under specific assumptions ^26^, but cannot establish whether an individual assignment is distinguishable from competing isobaric peptides. Although validation tools such as PepQuery and PepQuery2 compare non-canonical candidates against reference peptides on specified spectra ^27, 28^, reference competition alone does not systematically evaluate alternative non-canonical constructs, such as distinct junction arrangements or novel donor-fragment combinations. Synthetic peptide standards offer direct experimental verification ^12, 29, 30^, but require sequence selection prior to synthesis. Consequently, there remains a critical need for methods that explicitly evaluate all competing sequence explanations relevant to a non-canonical assignment.

To address this need, we developed TransPepMS, a software tool designed to validate non-canonical peptides by benchmarking candidate spectra against every isobaric alternative, including non-templated sequences. TransPepMS operates as a scalable pipeline on high-performance computing clusters and features a lightweight browser interface for rapid single-spectrum checks. We calibrated TransPepMS using simulated sequence pairs, evaluated its performance using same-mass entrapment controls on acquired spectra, and applied it to published cancer fusion peptides ^15, 31^, a contested spliced-peptide catalog ^5, 32^, and hybrid insulin peptides ^16^. TransPepMS provides sequence-level validation against explicit competitors alongside empirical false-acceptance rates, establishing a robust framework for biological interpretation and candidate prioritization.

## Results

### Overview of TransPepMS

TransPepMS evaluates candidate non-canonical peptides by assessing whether observed fragment ions uniquely distinguish them from competing sequence explanations (Fig. 1). Below, we describe how the tool selects these alternatives, compares their spectral support, and calibrates decision thresholds using benchmark spectra of known sequence.

**Figure 1.**
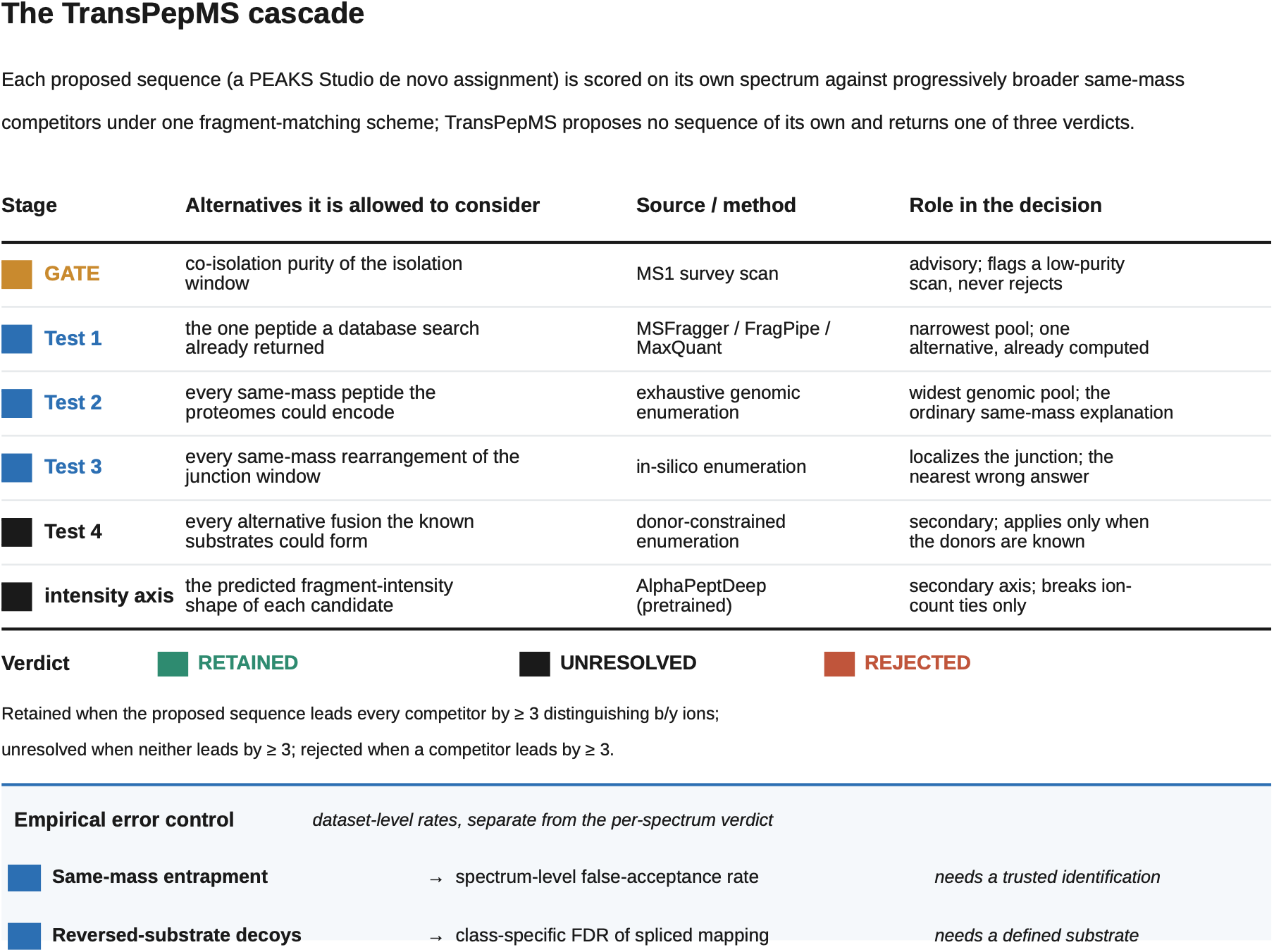
Spectral hypothesis competition. (a) A proposed sequence and its assigned spectrum, supplied by other software, are competed against each enumerated alternative sequence hypothesis of the same precursor mass, the proposed non-genomic sequence, a same-mass genome-encoded peptide, a same-mass rearrangement of the junction, and a same-mass alternative hybrid, each scored on that spectrum under one matching scheme. The verdict is retained (the proposed sequence is favored over every tested competitor by at least three distinguishing ions), unresolved (neither side is favored by three or more), or rejected (a competitor is favored over the proposed sequence by at least three). Abstention, not false acceptance, is the designed failure mode. (b) One co- isolation gate and four independent competitor pools, each a different class of same- mass alternative: the peptide a database search already returns (Test 1), every same- mass peptide the proteomes could encode, unmodified and then broadened to common modifications and known human variants (Test 2), every same-mass rearrangement of the junction window (Test 3), and every alternative hybrid the protein donors could form (Test 4), with predicted fragment intensity applied as a second scoring axis only where the ion count does not resolve the comparison within Test 4. Each applicable test is computed for a spectrum; the tests are independent. (c) The evaluation roadmap, in rising difficulty: simulated spectra where the truth is known by construction (Fig. 2), engineered impostors built to be rejected (Figure 6), transfer to an independent laboratory’s fusion- junction peptides (the GF-NEO dataset, PXD063253 ^15^; Figure 3), re-adjudication of a published spliced immunopeptidome (Fig.4), and the insulin-Spike peptides ^16^ that motivated the tool (Fig. 5). The dataset-by-component coverage is given in Supplementary Fig. S7.

The workflow begins with a precursor-purity assessment that flags substantial co-isolation without altering the final sequence verdict (Fig. 1; Supplementary Table ST1). TransPepMS then evaluates the proposed assignment through three primary tests. First, it compares the proposed assignment against the peptide an independent search engine assigned to the same scan (Test 1); second, it competes it against all explicitly enumerated isobaric, same-mass genome-encoded peptides (Test 2); and third, it evaluates the proposed junction against isobaric sequence rearrangements (Test 3).

When the input source proteins in a reaction mixture are known, an optional fourth test compares the candidate sequence against alternative hybrid products. This step provides secondary structural support without overriding the primary classification decision. Full procedures for generating candidate competitors are detailed in the **Methods** Section.

For direct ion comparisons, spectral support is quantified using the *distinguishing-ion margin*: the number of matched b/y ions unique to the proposed sequence minus those unique to its top-supported competitor. TransPepMS retains the proposed sequence if it passes every applicable primary test (Tests 1-3), rejects it if a competitor satisfies the rejection criteria in any primary test, and otherwise classifies the assignment as unresolved (Fig. 1).

The specific rejection rule applied depends on how the competing sequences are derived. For Test 1, TransPepMS compares the proposed peptide against the alternative sequence an independent search engine assigned to the exact same scan (e.g., MSFragger/FragPipe psm.tsv or MaxQuant), and rejects the proposed peptide if that independent assignment explains the spectrum equally well or better. For Tests 2 and 3, it enumerates all possible isobaric sequences encoded by the reference proteomes (Test 2) or all possible junction rearrangements (Test 3), regardless of whether an engine previously detected them; because these competitors are theoretical constructs rather than observed hits, a candidate is rejected only if a constructed competitor outperforms it by at least three distinguishing ions. If the enumeration yields no isobaric alternatives, the assignment is classified as mass-unique.

Following classification, TransPepMS estimates empirical false-acceptance rates using dataset-matched controls. *Same-mass entrapment* estimates how frequently an incorrect isobaric sequence is erroneously retained on acquired spectra, providing an error metric applicable to any dataset (including open-proteome discovery where candidate substrate sets are undefined). When source substrates are known, a *reversed-sequence substrate decoy* control additionally estimates the class-specific false discovery rate of mapping sequences to their proposed origin proteins. These two control strategies target distinct sources of error and are reported independently.

### Calibration of Distinguishing-Ion Thresholds Using Simulated Isobaric Pairs

Two peptides of distinct sequences can share an identical precursor mass; when they do, high-precision precursor mass measurements alone cannot distinguish them, leaving MS/MS fragment ion patterns as the sole diagnostic evidence. Selecting the correct sequence from an isobaric pair requires establishing a distinguishing-ion margin: the minimal excess of matched fragment ions by which the true sequence must lead its competitor to be confidently retained rather than left unresolved or rejected. Calibrating this margin requires prior knowledge of the true sequence for every spectrum so that false-retention events can be rigorously quantified. Experimental MS/MS spectra from complex biological samples cannot fulfill this requirement because the true sequence is the entity being inferred. To overcome this, we constructed a synthetic collection of 518,400 simulated isobaric peptide pairs, for which the generating sequence was recorded as the true sequence.

For each isobaric pair (A and B), we generated a simulated MS/MS spectrum from sequence A (the known correct sequence) and evaluated two distinct comparison arms against this spectrum: a correct-candidate arm, evaluating how sequence A scored relative to competitor B, and an incorrect-candidate arm, evaluating how sequence B (the false candidate) scored relative to A.

In the incorrect-candidate arm, false-retention events decreased steeply as the required margin increased: from 1,862 false retentions at a one-ion margin (3.6 x 10^-3^) to 27 at a two-ion margin (5.2 x 10^-5^), and dropping to zero at a three-ion margin or greater (Fig. 2a). Based on these empirical distributions, we set the retention threshold at three distinguishing ions, the smallest margin at which no incorrect candidate was retained across all 518,400 tests (one-sided 95% upper confidence bound: 5.8 x 10^-6^).

**Figure 2.**
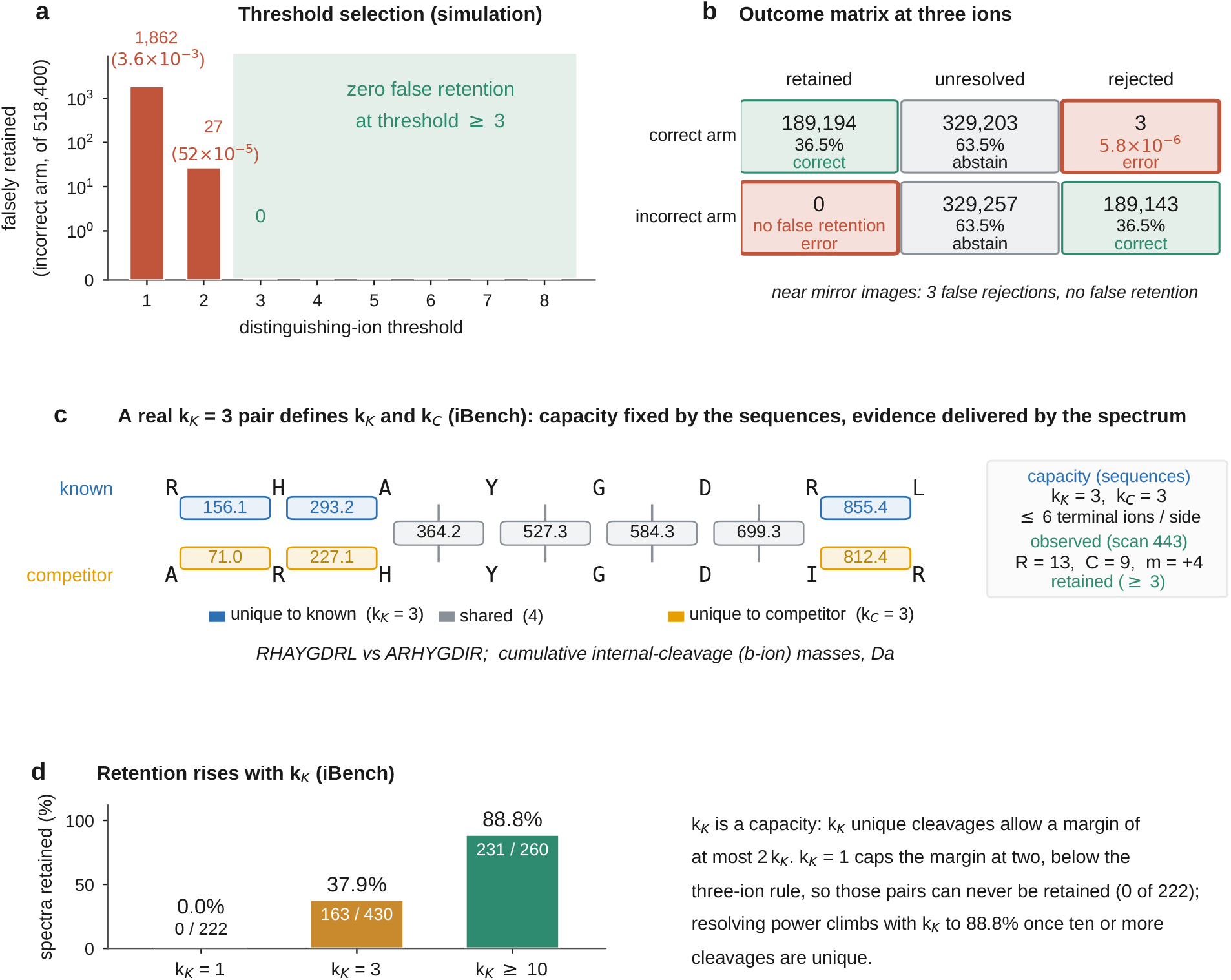
Selecting the three-ion rule. Panels a and b are the simulation, two matched arms of 518,400 comparisons each with the generating sequence known by construction; panels c and d are the iBench acquired benchmark (PXD044451), 4,374 spectra of 1,183 known peptides each competed against every same-mass peptide the six reference proteomes encode. kK, the reference-specific count of distinguishing fragmentations, is the number of cumulative b-ion (prefix) masses unique to the known benchmark peptide. kC, the competitor-specific count, is the number unique to its closest competitor. Each caps the distinguishing terminal-ion evidence at 2kK and 2kC, so a pair with too few distinguishing fragmentations cannot reach the three-ion margin at any spectrum quality. (a) Threshold selection: in the incorrect-candidate arm (518,400 comparisons) the number falsely retained falls from 1,862 at a one-ion margin (3.6 × 10⁻³) to 27 at two (5.2 × 10⁻⁵) to zero at three ions and above, so three is the least stringent margin with no false retention. (b) Outcome matrix at three ions: the two arms give symmetric outcomes (36.5% retained vs rejected and 63.5% unresolved on each side), with correct candidates 189,194 retained / 329,203 unresolved / 3 rejected and incorrect candidates 0 retained / 329,257 unresolved / 189,143 rejected; the only errors are three false rejections and no false retention. (c) A real kK = 3 pair (RHAYGDRL vs ARHYGDIR, scan 443) defines kK and kC: every backbone fragmentation and its cumulative b-ion mass, colored gray where the two sequences share the mass, blue where it is unique to the known peptide (kK = 3), and orange where it is unique to the competitor (kC = 3). These counts are a capacity fixed by the sequences (at most six distinguishing terminal ions per side), separate from the evidence the spectrum delivers (R = 13 and C = 9 matched ions, margin m = +4, retained). (d) Across the 1,911 spectra with an enumerable competitor the retained fraction rises with kK: 0/222 at kK = 1, 163/430 (37.9%) at kK = 3, and 231/260 (88.8%) at kK ≥ 10. The decision-rule schematic and the correct-arm retention across thresholds are in Supplementary Fig.S1; peptide length and per-substrate views in Supplementary Fig. S8.

Applying this three-ion threshold to the correct-candidate arm successfully retained 189,194 true assignments (36.5%), left 329,203 unresolved (63.5%), and rejected 3 (Fig. 2b). Increasing the threshold to four ions similarly yielded no false retentions but reduced correct candidate retention to 28.7%.

We adopted the three-ion margin for all subsequent analyses. The trade-off of this conservative threshold is coverage rather than accuracy: unassigned correct candidates are predominantly classified as unresolved rather than misassigned to the wrong sequence. Thus, TransPepMS abstains when spectral evidence is ambiguous rather than making a false call.

### Theoretical Ceilings on Sequence Discrimination

At the three-ion margin, most correct assignments were left unresolved rather than retained. Whether this reflects a weakness of the method or a property of the data it is given depends entirely on why a call is left unresolved, and there are two possibilities. Either the spectrum recorded too few of the distinguishing ions that the sequence pair is capable of producing, in which case a higher-quality spectrum would resolve it, or the pair cannot produce three distinguishing ions at any spectral quality, because the two sequences are too similar, in which case no spectrum on any instrument can resolve it. The first is a limitation of the data; the second is fixed by the sequences themselves, before any spectrum is acquired. The rest of this section establishes that the intrinsic limit is what dominates, and that it can be computed in advance.

That intrinsic limit follows directly from the primary structures. Once two candidate sequences are specified, the number of backbone fragment masses that differ between them is fixed, and each differing mass can yield at most two distinguishing ions, a b-ion and its complementary y-ion. We term each such position a distinguishing fragmentation and denote its count k, writing kK for the known peptide and kC for its closest competitor. The known peptide can therefore lead its competitor by at most 2kK distinguishing ions. When kK = 1 the largest possible lead is two ions, short of the three-ion threshold, so no spectrum from that pair can ever retain the correct assignment; when kK = 3, as for the pair RHAYGDRL and its closest competitor ARHYGDIR, up to six distinguishing ions are available, and in the acquired spectrum RHAYGDRL led by four ions (13 versus 9 matched) and was retained (Fig. 2c).

To test whether this ceiling, rather than spectral quality alone, governs the outcome, we used iBench, a benchmark in which the true sequence behind every spectrum is known because the peptides are produced by *in vitro* digestion of synthetic substrates with the human 20S proteasome (4,374 spectra from 1,183 peptides; PXD044451) ^32^. Each known peptide was competed against every same-mass peptide encoded by six reference proteomes. Of the 4,374 spectra, 1,911 had at least one same-mass competitor and could therefore be scored; among these, retention rose monotonically with the ceiling: 0 of 222 pairs were retained at kK = 1, 163 of 430 (37.9%) at kK = 3, and 231 of 260 (88.8%) at kK ≥ 10 (Fig. 2d). Retention is therefore set by how many distinguishing ions the sequences can produce, not by random variation in spectral quality.

Abstention is therefore not a single failure mode. An unresolved call means one of two things: either a spectrum too sparse for a pair that is intrinsically resolvable, which deeper acquisition or a synthetic standard can address, or a pair that cannot reach the three-ion margin at any spectral quality, which no spectrum can resolve and which instead requires orthogonal evidence such as retention time or predicted fragment intensity. Because the ceiling is computed from the two candidate sequences alone, TransPepMS reports which of these applies before any further data are acquired, separating a solvable data-quality problem from a fundamental limit of fragment-based sequencing.

### Validation of the Three-Ion Rule on Independent Reference Spectra

Up to this point, the three-ion rule had been calibrated solely on simulated or curated spectra, where control sequences were fixed by design or established by benchmark sets. Because decision thresholds fit to idealized datasets may not generalize to raw experimental data, we evaluated the unadjusted rule on independent, real-world acquired spectra whose correct sequences were established independently. Specifically, we applied the three-ion margin requirement without modification to 200 high-confidence canonical peptide-spectrum matches (PSMs) from a HeLa cell line dataset (PXD000508)^33^. As a negative control, we tested 46 incorrect *de novo* sequence predictions derived from the same spectra as known-false assignments.

The three-ion rule correctly retained 188 of the 200 true-positive assignments, left 11 unresolved, and rejected one. The single rejection resulted from a positional misassignment of methionine oxidation. Re-evaluating the spectrum with the corrected oxidation site yielded an 11-fragment match with a four-ion lead over its nearest competitor. Crucially, none of the 46 incorrect *de novo* predictions were retained; all were classified as unresolved. No incorrect de novo prediction was retained among the 46 negative controls (one-sided 95% upper confidence limit 6.3%).

Thus, on experimental spectra with known answers, the three-ion rule successfully retained true-positive sequences while completely excluding known-incorrect assignments. Its primary failure mode was abstention rather than misclassification.

### TransPepMS detects and adjudicates independently reported fusion neoantigens

The previous tests relied on peptides of known established identity. We next evaluated whether TransPepMS could independently adjudicate non-canonical assignments generated outside its development framework. We analyzed the GF-NEO dataset (gene-fusion neoantigens; PXD063253), an HLA class I immunopeptidome from pediatric cancer samples in which candidate sequences and their genomic breakpoints had been identified in an independent study ^15^. The six target neoantigens span the fusion junctions of ETV6-RUNX1 or CBFB-MYH11, two driver gene fusions in acute leukemia resulting from chromosomal translocations that generate tumor-specific junction sequences. For each reported neoantigen, we evaluated spectrum detection, junction localization (Test 3), and same-mass competition (Test 2).

All six reported sequences were detected, yielding matched b/y-ion coverage ranging from 7 to 15 ions (Fig. 3a; Supplementary Table ST6). Same-mass competition then adjudicated the recovery-threshold-positive peptides: across 16 PSMs, 3 were retained, 11 remained unresolved, and 2 were rejected (Fig. 3b). Junction localization provided an orthogonal, position-level readout: five sequences localized their reported junctions in fusion-expressing samples (junction competitor ratios 0 to 0.008, well below the 0.05 threshold; Fig. 3c).

**Figure 3.**
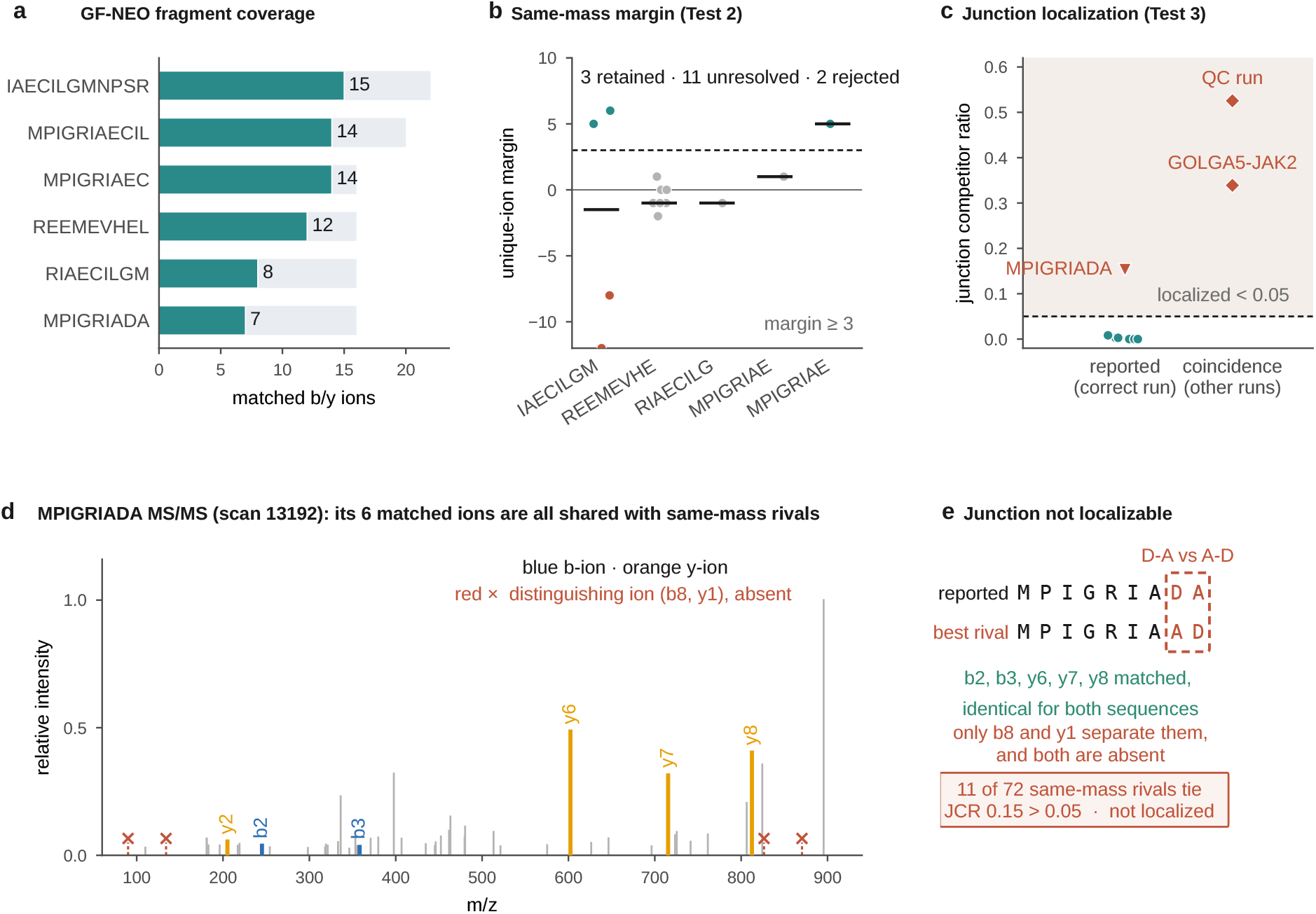
External validation on six independently reported fusion-junction hybrid peptides from two fusion systems (GF-NEO, PXD063253). The six reported junction peptides of ETV6-RUNX1 and CBFB-MYH11 were taken as candidates and searched in the deposited peak lists; the authors’ assignments were used only for comparison. (a) GF-NEO fragment coverage: matched b/y ions at detection against the eight-ion recovery threshold; all six peptides were detected, five were recovery-threshold positive, and MPIGRIADA was detected at seven matched b/y ions (below the threshold, in amber). (b) Same-mass competition (Test 2): the best same-mass distinguishing-ion margin per peptide (per-PSM medians shown); across the 16 adjudicated peptide-spectrum matches, 3 were retained, 11 unresolved, and 2 rejected at the three-ion margin, both rejections being the IAECILGMNPSR quality-control mass-coincidence. MPIGRIADA was not carried into same-mass competition. (c) Junction localization (Test 3): the five recovery-threshold-positive peptides localized at a junction competitor ratio of 0.05 or below in the sample expressing their fusion, whereas MPIGRIADA did not localize (junction competitor ratio 0.15; amber triangle). The two IAECILGMNPSR mass- coincidences in unrelated runs (a GOLGA5-JAK2 run and a quality-control run) were not identified (junction competitor ratios 0.34 and 0.53) and serve as negative controls. (d) MPIGRIADA tandem mass spectrum (scan 13192): the six matched fragment ions (blue b-ions, orange y-ions) are all shared with its same-mass rivals, and the two ions that would distinguish the reported sequence, b8 and y1, are absent (red). (e) Why the MPIGRIADA junction cannot be localized: the reported sequence (ending D-A) and its best same-mass rival (ending A-D) are separated only at the terminal junction; b2, b3, y6, y7 and y8 are identical for both sequences, only b8 and y1 distinguish them, and both are absent, so 11 of 72 enumerated same-mass rearrangements tie (junction competitor ratio 0.15 > 0.05; not localized). Six-proteome same-mass competition, 17,210 to 30,999 competitors per spectrum, as for the other datasets.

In contrast, the peptide MPIGRIADA failed junction localization. Its ETV6-RUNX1 junction occurs between the seventh and eighth residues (MPIGRIA\DA), meaning only the C-terminal DA motif originates from RUNX1. The two fragment ions required to span this junction (b8 and y1) were both absent from the spectrum (Fig. 3d). The six detected fragment ions are shared between the reported sequence and its positional rearrangements; consequently, 11 of the 72 enumerated same-mass candidate rearrangements matched the spectrum with equal fidelity (junction competitor ratio: 0.15; Fig. 3e). Deprived of its diagnostic ions crossing the terminal junction, the localization module classified this assignment as unresolved based on its own spectral evidence.

The peptide IAECILGMNPSR demonstrated the specificity of the localization filter. It successfully localized in the fusion-expressing sample, yet the same sequence also matched, by mass coincidence, in two fusion-negative controls (a GOLGA5-JAK2 run and a quality-control run). In both negative controls, the junction failed to localize (junction competitor ratios: 0.34 and 0.53). Thus, localization triggered exclusively in the presence of the true gene fusion rather than on coincidental same-mass matches.

In summary, detection, same-mass competition, and junction localization provided distinct diagnostic parameters for each assignment. While all six sequences were detected, junction localization selectively validated five, and same-mass competition classified the localized spectra into retained, unresolved, and rejected categories. Although restricted to a modest candidate set, these results demonstrate that TransPepMS reliably adjudicates non-canonical assignments derived from independent computational pipelines.

### Few Contested Spliced Assignments Survive Isobaric Competition

While the fusion analysis evaluated a small set of defined junction peptides, we next applied TransPepMS to a large, previously contested dataset (PXD000394 ^5^) containing 1,154 unique human HLA class I immunopeptidome sequences originally attributed to proteasome splicing. Because proteasome splicing links non-adjacent fragments, spliced peptides are not contiguously encoded in the genome. Independent reanalyses subsequently challenged these identifications, proposing that standard genome-encoded (canonical) isobaric peptides could explain the observed spectra ^17, 18^. We tested each reported sequence on its assigned scan against all isobaric canonical alternatives (Test 2).

The advisory precursor-purity gate first flagged 216 of the 1,154 assignments (18.7%) for potential co-isolation (Fig. 4a). Isobaric competition (Test 2) then retained only 53 of the 1,154 reported assignments (4.6%), left 718 unresolved (62.2%), and rejected 383 (33.2%) (Fig. 4b; Supplementary Table ST7), while junction localization (Test 3) supported 638 of 1,125 evaluable junctions (56.7%; Fig. 4c). These outcomes are robust to the decision threshold and to held-out validation: when the distinguishing-ion threshold was selected on a held-out split of the 1,036,800-case simulation it retained zero false positives on the test arm (0 of 24,586; 95% Wilson upper bound 0.016%), a stricter four-ion threshold leaves 22 retained, and on the decisive cross-engine calls the orthogonal engines named the same genome-encoded competitor in 809 of 861 (94.0%). Prior to applying the three-ion threshold, the reported spliced sequence led its top canonical competitor in 245 cases (21.2%), tied in 160 (13.9%), and trailed a canonical sequence in 749 (64.9%).

**Figure 4.**
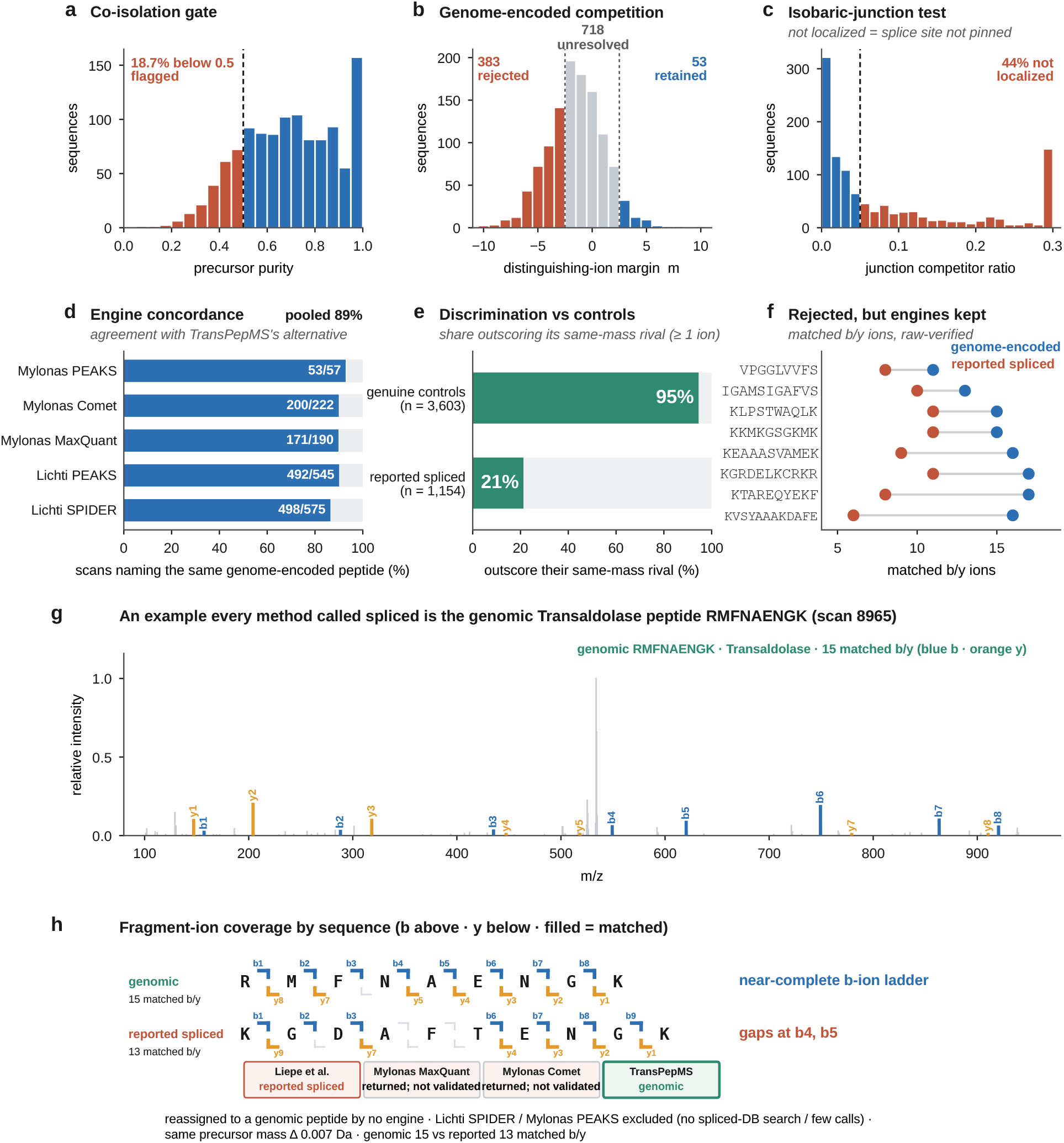
Genomic alternatives explain most reported peptides in a published spliced immunopeptidome (PXD000394). The 1,154 unique originally reported spliced sequences, adjudicated at the scans the authors assigned. (a) Co-isolation gate: precursor purity per sequence, with sequences below the 0.5 advisory threshold flagged for review. (b) Same-mass genomic competition: the distinguishing-ion margin against the enumerated same-mass genome-encoded competitors gives 53 retained, 718 unresolved, and 383 rejected on which sequence leads at all, the reported spliced sequence leads for 245, ties for 160, and a genome-encoded competitor leads for 749. (c) Isobaric-junction test: the junction competitor ratio among the 1,125 sequences with a defined junction indicates 638 junctions are identified below a 0.05 threshold. (d) Engine concordance: among the scans where TransPepMS rejected the reported sequence and an independent search engine also assigned that scan to a same-mass genome-encoded peptide per engine (Mylonas PEAKS 53/57 ^18^, Comet 200/222 ^34^, MaxQuant 171/190; Lichti PEAKS 492/545 ^19^, SPIDER 498/575 ^15^) and pooled (1,414/1,589, 89.0%). Each denominator is that engine’s count of comparable scans, not its total genomic reassignments; pooled comparisons are not independent, as one scan can contribute to more than one engine. (e) Discrimination against controls: the reported spliced sequence is favored by distinguishing ions for 21.2% of cases after re-evaluation. (f) Rejected, but engines kept: representative sequences that a search engine retained yet a same-mass genome-encoded peptide outscored on matched b and y ions. Purity, junction, and full per-sequence counts are in Supplementary Table ST2.

The genome-encoded alternatives identified by Test 2 aligned closely with output from independent search engines. Comparing scans where Test 2 rejected or left the reported sequence unresolved and an independent engine assigned an isobaric canonical peptide, TransPepMS matched the independent assignment in 1,414 of 1,589 eligible comparisons (89.0%; Fig. 4d; Supplementary Table ST8). Agreement by individual engine was 53/57 for Mylonas PEAKS ^18^, 200/222 for Comet ^34^, 171/190 for MaxQuant, 492/545 for Lichti PEAKS ^19^, and 498/575 for SPIDER ^15^. While not an independent validation, since individual scans can enter multiple comparisons and all engines analyze the same spectra, this high concordance demonstrates that explicit isobaric competition consistently identifies the same alternative sequence explanations as dedicated database search pipelines.

We also benchmarked discrimination on sequences with established ground truth. A positive distinguishing-ion margin separated 94.5% of the dataset’s 3,603 trusted canonical identifications from their isobaric alternatives, compared to only 21.2% of the reported spliced sequences (Fig. 4e). These fractions describe which sequence led before the decision threshold; they are not retention rates. Representative rejected assignments and their top canonical competitors are detailed in Fig. 4f and Supplementary Table ST3, with complete outcome distributions, margin metrics, and engine agreement summarized in Supplementary Fig. S5.

In summary, under explicit isobaric competition (Test 2), fewer than 1 in 10 contested assignments met the retention criterion (53/1,154). The vast majority were either left unresolved due to insufficient distinguishing fragment ions or rejected outright in favor of a canonical genome-encoded peptide. Because these assignments were previously disputed, these 53 sequences carry explicit isobaric support distinct from the larger pool easily explained by canonical sequences. Nine of the 53 (17%) had precursor co-isolation purity below the 0.5 advisory flag (lowest MIFFVSGKI 0.25, KIQDKTLTGK 0.29, IYVGLKGML 0.44); the co-isolation gate marks these for review but, being advisory, does not reject them.

### Isobaric Competition Supports Insulin-Spike Hybrids

Because the donor proteins in the insulin-spike reactions are defined, the full TransPepMS cascade, including donor-hybrid competition (Test 4), could be applied. These hybrid peptides are formed when cathepsin S joins insulin and spike protein fragments via reverse proteolysis ^16^. We evaluated 145 sequences: 99 reported insulin- spike hybrids ^16^ and 46 canonical genome-encoded peptides inserted as negative controls ("planted false hybrids"). TransPepMS retained 91 of the 99 hybrids, rejected none, and correctly excluded all 46 planted controls (Fig. 5a). The precursor-purity advisory gate flagged 5 of the 99 hybrid spectra while leaving 94 above the purity threshold of 0.5 (Fig. 5b).

**Figure 5.**
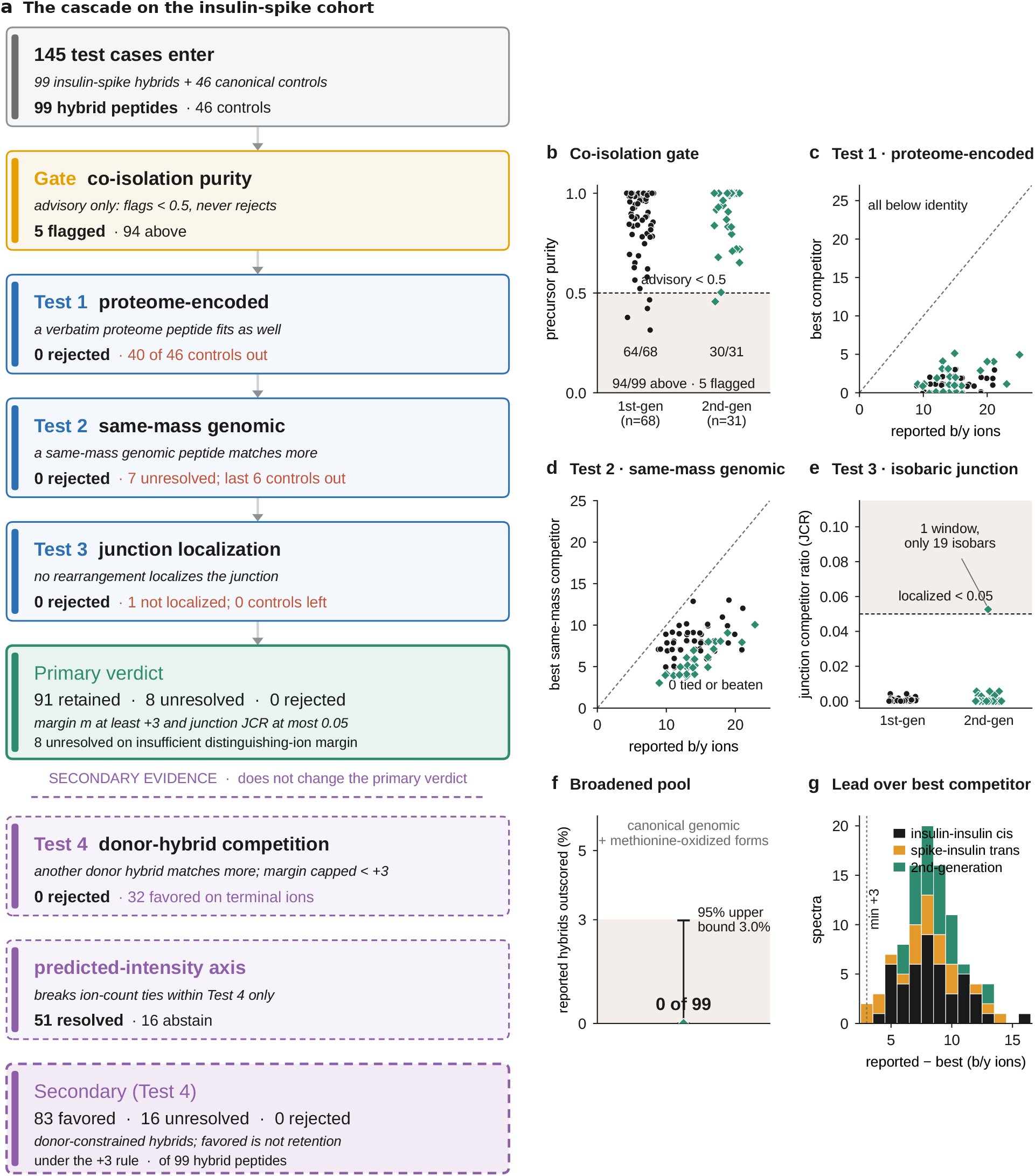
The cascade on the insulin-spike cohort. (a) Cascade summary: 145 test cases enter (99 insulin-spike hybrid spectra, representing 98 unique sequences with the hybrid LGAENSVASHLVE observed in two spectra, and 46 canonical controls). The advisory co-isolation gate flags 5 and rejects none; Test 1 (proteome-encoded) removes 40 of 46 controls; Test 2 (same-mass genomic) removes the remaining 6 controls; Test 3 (junction localization) leaves 1 hybrid unlocalized. The primary verdict (same-mass margin ≥ 3 with junction competitor ratio ≤ 0.05) is 91 retained, 8 unresolved (7 on the ion-count margin, 1 on localization) and 0 rejected. Secondary evidence (Test 4 donor- hybrid competition and the predicted-intensity axis) does not change the primary verdict; Test 4 favors 83 of 99, leaves 16 unresolved and rejects none. (b) Co-isolation purity per peptide for first-generation (n = 68) and second-generation (n = 31) hybrids; 94 of 99 lie above the 0.5 advisory threshold and 5 are flagged. (c) Test 1: reported b/y ions versus the best proteome-encoded competitor, all below the identity line. (d) Test 2: reported b/y ions versus the best same-mass genomic competitor, none tied or beaten. (e) Test 3: junction competitor ratio by generation, with reported junctions localizing below 0.05. (f) Broadened same-mass competitor pool (canonical genomic and methionine-oxidized forms): no competitor outscored a reported hybrid on any of the 99 spectra (95% upper bound 3.0%). (g) Distribution of the reported sequence’s lead over its best competitor (reported − best competitor b/y ions) for insulin-insulin cis, spike-insulin trans and second- generation hybrids; the dashed line marks the +3 margin.

All 46 planted controls were eliminated by the canonical proteome tests: 40 matched a canonical proteome peptide directly (Test 1), and 6 were outscored by a superior isobaric canonical match (Test 2). In contrast, across the 99 reported hybrids, no candidate sequence was tied or outscored by a proteome-encoded competitor (Test 1; Fig. 5c) or an isobaric canonical competitor (Test 2; Fig. 5d). Furthermore, all proposed junctions localized below the 0.05 competitor ratio except one (Test 3; Fig. 5e). Under the primary rule (a same-mass distinguishing-ion margin of at least three, combined with junction localization, Test 3), 91 hybrids were retained, 8 were unresolved, and zero were rejected (Fig. 5a). Because all negative controls were eliminated and no reported hybrid was rejected, this reflects robust discrimination rather than unselective retention. Unresolved cases stemmed from insufficient distinguishing spectral evidence rather than superior competing sequences.

This performance persisted when expanding the competitor pool. Each of the 99 reported hybrids was evaluated against a broadened same-mass competitor pool of canonical genomic peptides and their methionine-oxidized forms. No competitor outscored a reported hybrid on any of the 99 spectra (95% upper confidence bound: 3.0%; Fig. 5f). Secondary analyses reinforced these findings: exhaustive donor-hybrid competition (Test 4), which scores each reported hybrid against every same-mass peptide the two donor proteins can assemble from up to three fragments, up to billions of configurations per spectrum (Methods), favored the reported sequence in 83 of 99 cases (16 unresolved, 0 rejected), while the predicted-intensity module resolved 51 of the terminal-ion-unresolved comparisons (Fig. 5 a,g).

Only one sequence-defined edge case remained unresolved: SHLVEYTPKT, a *cis*- spliced product that differs from its exact isobar by a single backbone cleavage site. Because only two fragment ions distinguish this pair (Supplementary Tables ST4, ST5), the margin falls below the three-ion threshold, triggering an automated "unresolved" verdict despite independent verification by a synthetic standard ^16^.

An independent survival analysis across a broader cohort of 108 hybrids (75 first-generation, 33 second-generation) further confirmed these assignments. Of these, 105 were mass-unique against a modification-aware competitor database, and the remaining 3 were resolved by junction localization (Test 3), yielding Clopper-Pearson survival rates of 75/75 (95% CI: 95.2-100%) and 33/33 (95% CI: 89.4-100%) ^16^. That an independent criterion on an expanded cohort yielded identical conclusions confirms that support for these hybrids is robust and independent of specific cohort filtering or decision thresholds. For most of these assignments no same-mass genome-encoded peptide existed to compete, so they carried no isobaric ambiguity; the few that did were resolved by junction placement rather than ion counting.

### Direct Comparison with PepQuery: Isobaric Competition Overturns Retained Calls and Resolves Hybrid Rivals

To benchmark TransPepMS against single-spectrum validation tools, we compared its performance to PepQuery, the primary reference-competition tool in the field ^27, 28^. While PepQuery evaluates whether a spectrum matches a candidate better than the best reference peptide in a canonical database, TransPepMS competes candidates against explicitly enumerated isobaric alternatives, including reordered junctions and non-canonical donor hybrids. Comparing both tools on the contested Liepe dataset (PXD000394) ^5^ reveals how explicit isobaric competition alters sequence calls.

We next set the two tools against each other on the same 1,154 reported spliced peptides. PepQuery2 scored 200 of them as confident; of those 200, TransPepMS retained 33 (16.5%), left 134 (67.0%) unresolved, and rejected 33 (16.5%) (Fig. 6a). The disagreement is asymmetric (McNemar χ² = 114.0, P < 0.001). These are the PepQuery- confident share of the full re-adjudication reported above: the 33 retained and 33 rejected come out of the 53 peptides TransPepMS retained and the 383 it rejected across all 1,154, with the remaining 20 of the 53 retained by TransPepMS but not scored confident by PepQuery2. The 33 are the decisive disagreement: PepQuery2 confidently validated each of these spliced calls, yet TransPepMS found a same-mass genome-encoded peptide that explained the spectrum by at least three more distinguishing ions. Scan 30840 is one such case. PepQuery2 marked the spliced sequence GPPGSVILY as confident, but the same-mass genome-encoded peptide ILGPPGSVY (from ubiquitin-conjugating enzyme E2, UBE2E1/2/3) led the spliced sequence by four distinguishing b/y ions on the same spectrum (distinguishing-ion margin −4; Fig. 6b), uniquely explaining the two most intense peaks (y6 and y7), which the spliced sequence cannot produce. Per-spectrum same-mass competition identified this canonical alternative and reversed the call.

**Figure 6.**
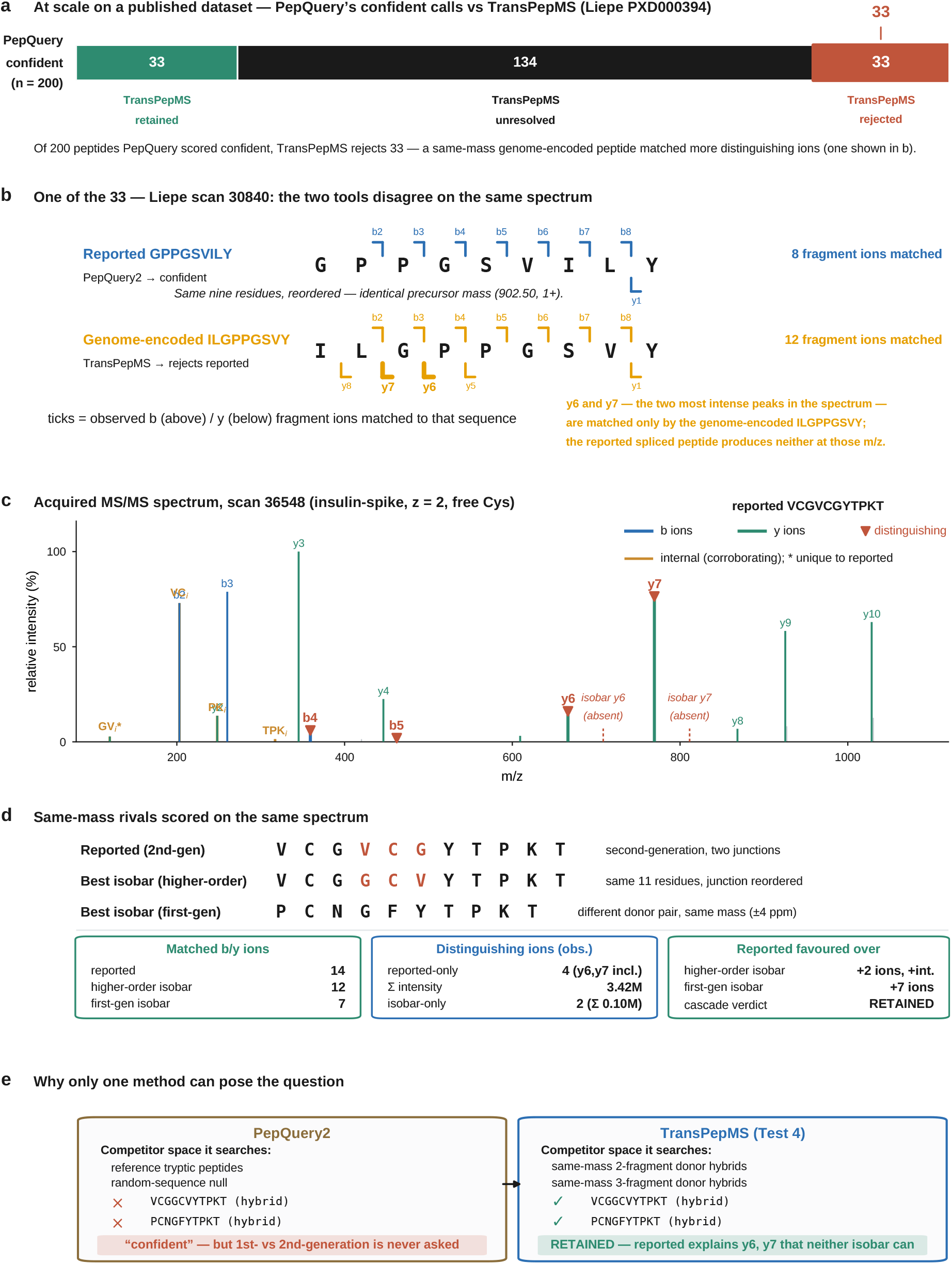
Explicit isobaric competition overturns PepQuery-confident calls and resolves hybrid rivals that reference competition cannot evaluate. (a) At scale on the contested Liepe dataset (PXD000394): of 200 peptides scored confident by PepQuery2, TransPepMS retained 33, left 134 unresolved, and rejected 33, each rejection because a same-mass genome-encoded peptide matched more distinguishing ions. (b) One rejected example (Liepe scan 30840): PepQuery2 scored the reported spliced peptide GPPGSVILY confident, matching 8 fragment ions in its own search; on the same spectrum the same-mass genome-encoded peptide ILGPPGSVY, a reordering of the same nine residues at an identical precursor mass (902.50, 1+), is favoured by TransPepMS’s distinguishing-ion margin (−4), uniquely explaining the two most intense peaks, y6 and y7, which the spliced sequence cannot produce. (c) Acquired tandem mass spectrum of the two-junction insulin-spike hybrid VCGVCGYTPKT (scan 36548; z = 2, free cysteine), showing matched b-ions (blue), y-ions (green), corroborating internal ions (orange), and the distinguishing ions (red). (d) The same spectrum scored against its same-mass rivals: the reported second-generation hybrid matched 14 b/y ions, versus 11 for the best higher-order isobar (VCGGCVYTPKT, the junction reordered) and 6 for the best first-generation isobar from a different donor pair (PCNGFYTPKT, within 4 ppm); four observed ions were unique to the reported sequence, including the dominant y6 and y7, and the cascade retained it. (e) Why only TransPepMS can pose the question: PepQuery2 searches reference tryptic peptides and a random-sequence null and cannot evaluate the unindexed same-mass hybrids, so first- versus second-generation hybrid origin is never tested, whereas TransPepMS donor-hybrid competition (Test 4) enumerates both and favours the reported sequence because it alone explains y6 and y7.

PepQuery2 validates candidates against reference proteomes and randomized sequences, but cannot evaluate unindexed isobaric hybrids. This limitation is particularly critical for complex non-canonical peptides, such as the two-junction insulin-spike hybrid VCGVCGYTPKT (scan 36548; free cysteine, 2+ charge). Although PepQuery2 reported a confident match, its reference search could not evaluate two key isobaric hybrid rivals: VCGGCVYTPKT (a higher-order isobaric rearrangement) and PCNGFYTPKT (a first-generation hybrid from a different donor pair within 4 ppm). TransPepMS enumerated both rivals and evaluated all three candidates against the experimental spectrum (Fig. 6 c,d). The reported sequence matched 14 b and y ions, outscoring the higher-order isobar (11 matched ions) and the first-generation isobar (6 matched ions). Four observed fragment ions were unique to the reported sequence, including the predominant y6 and y7 peaks, while the higher-order isobar matched only one minor unique ion (Fig. 6 c,d). The resulting three-ion distinguishing margin over the higher-order isobar met the automated primary threshold, retaining the reported sequence, and the predicted- intensity axis independently corroborated the call. Because non-canonical isobaric hybrids are absent from canonical reference databases and random decoys, standard reference-competition searches cannot evaluate them or distinguish second-generation assignments from reordered sequence variants (Fig. 6e). TransPepMS donor-hybrid competition (Test 4) addresses this gap, favoring the reported sequence in 83 of the 99 tested hybrids (Fig. 5a).

### Same-Mass Entrapment Estimates False Acceptance on Acquired Spectra

To measure how often TransPepMS falsely retains an incorrect sequence, we used same-mass entrapment. For each acquired spectrum whose correct peptide is already trusted, we constructed a decoy by joining fragments of real proteome sequence into a deliberately incorrect peptide of the same precursor mass as the trusted peptide, verified neither to occur in the proteome nor to equal the true sequence. Each decoy, wrong by construction, was then scored against its real spectrum through the full cascade with the true peptide kept in the competitor pool; any retained decoy counts as a false acceptance, and scoring the trusted peptides the same way gives a paired positive control for how often a correct answer is kept. Because each decoy is mass-matched to a genuine peptide and tested on a genuine acquired spectrum, this measures false acceptance under realistic conditions rather than in simulation. Same-mass entrapment is one of two error-control strategies that suit different discovery regimes (Fig. 7a,b).

**Figure 7.**
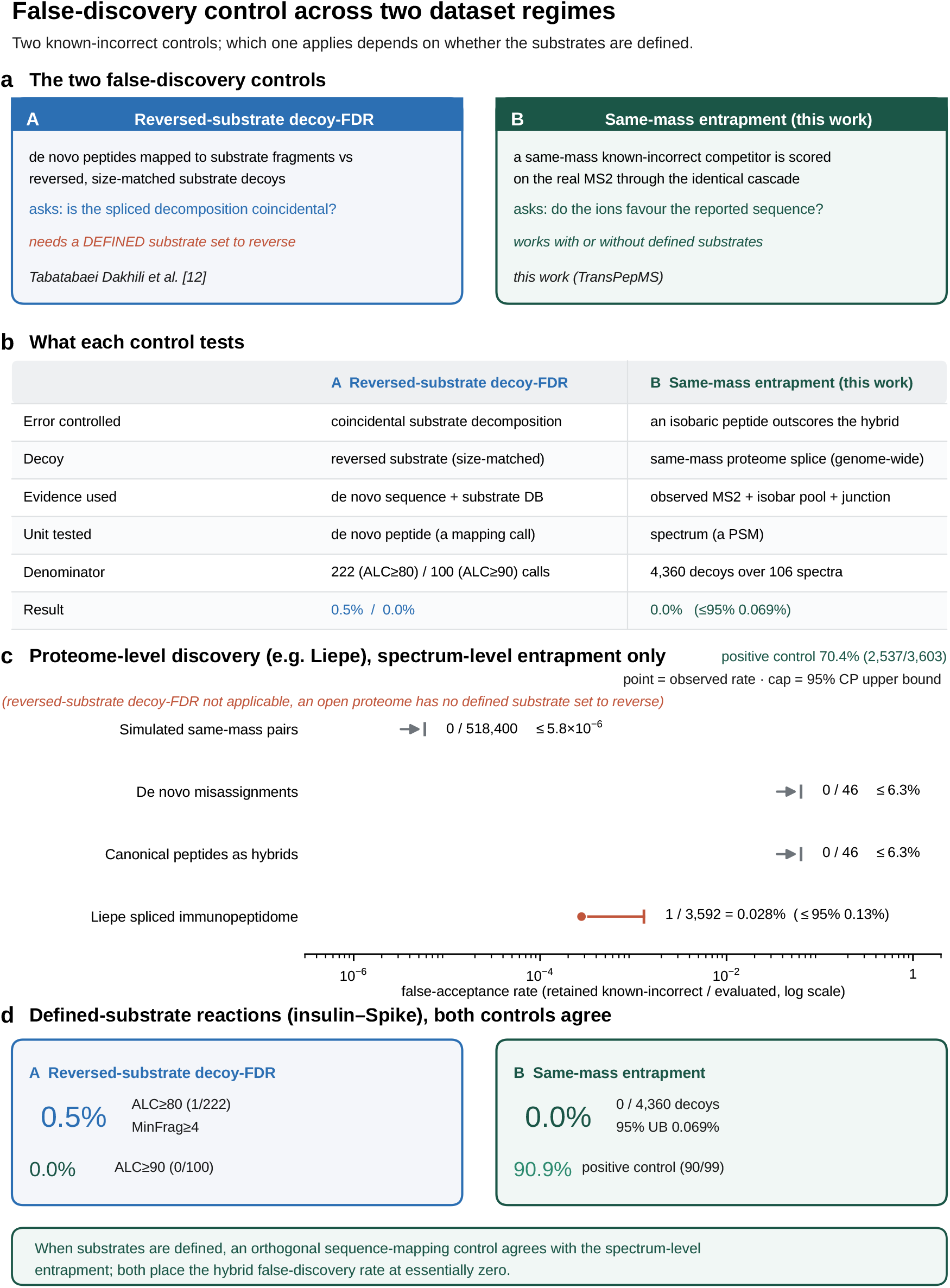
**Empirical error control across two discovery regimes**. (a) The two known- incorrect controls: a spectrum-level same-mass entrapment (this work), which scores a same-mass incorrect competitor on the real spectrum, and a sequence-mapping reversed-substrate decoy-FDR ^16^). (b) Axis-by-axis comparison of what each control tests. (c) Proteome-level discovery (e.g., ^5, 32^: only the spectrum-level entrapment applies, because an open proteome has no defined substrate set to reverse; no known-incorrect candidate was retained across the simulated, de novo and canonical-as-hybrid controls, and 1 of 3,592 was retained in the Liepe canonical set (0.028%; one-sided 95% Clopper- Pearson upper bounds shown), with a 70.4% positive control. (d) Defined-substrate reactions (insulin-Spike): both controls apply and agree, the reversed-substrate decoy- FDR (0.5% at ALC>=80, 1/222; 0.0% at ALC>=90, 0/100) and the same-mass entrapment (0 of 4,360, one-sided 95% upper bound 0.069%; 97.2% positive control, 106/109).

We first applied same-mass entrapment to the disputed PXD000394 immunopeptidome. Of 3,592 decoys built from its trusted canonical spectra, the cascade retained only 1 and rejected 3,591, a false-acceptance rate of 0.028% (one-sided 95% Clopper-Pearson upper bound 0.13%; Fig. 7c); the single survivor, LISPLKLKL, led the true sequence LISPIILKK by exactly three distinguishing ions, sitting at the retention threshold. In the paired positive control on the 3,603 trusted canonical identifications from which these decoys were built, the cascade retained 2,537 (70.4%) and rejected only 13 (0.4%), leaving 1,053 (29.2%) unresolved rather than misassigned (this positive control evaluates the same-mass competition criterion, Test 2, only; the Test-1 genome-encoded rejection is not applied to these genome-encoded controls), reflecting spectra with too few distinguishing ions rather than a competing sequence. The same decision rule applied to three further sets of known-incorrect sequences produced no false retentions at all: 518,400 simulated comparisons, 46 incorrect *de novo* identifications (PXD000508), and 46 canonical control spectra from the insulin-spike cohort (Fig. 7c).

### Error Control Depends on the Discovery Regime

Optimal error estimation depends directly on how candidate sequences are generated (Fig. 7 a,b). Same-mass entrapment evaluates spectrum-level false acceptance by competing an incorrect isobaric sequence against a spectrum with a trusted identity, keeping the true sequence in the competitor pool, and counting any retention of the entrapment sequence as a false acceptance. This metric is universally applicable whenever spectra with high-confidence identifications are available.

By contrast, defined-substrate experiments enable an additional error-control strategy. Reversed-substrate decoys reverse known substrate sequences and re-run the splice- mapping pipeline against them. Because a *de novo* sequence that decomposes into fragments of a reversed substrate can only do so by chance, the proportion of *de novo* reads mapping to decoy substrates estimates the false discovery rate (FDR) of the splice- mapping step ^16^. Crucially, these two control strategies evaluate distinct failure modes rather than two estimates of the same underlying metric.

For PXD000394, same-mass entrapment and the positive control described above provide the only applicable spectrum-level error estimates, as no defined substrate set exists to construct reversed-substrate decoys. For defined-substrate systems, such as the insulin-spike reactions, both controls can be applied. Reversed-substrate mapping yielded 1 decoy assignment among 222 spliced-callable peptides at an Average Local Confidence threshold of ALC ≥ 80% (0.5%) (FDR = 0.5%) and 0 decoy assignments among 100 peptides at ALC ≥ 90% Concurrently, same-mass entrapment retained 0 of 4,360 known-incorrect candidates (one-sided 95% upper confidence bound: 0.069%; Fig. 7d). In the corresponding positive control, 91 of 99 reported hybrid spectra (91.9%) satisfied the same-mass competition criterion (Test 2). Note that this metric evaluates Test 2 alone, whereas the deduplicated primary pipeline also enforced junction localization via Test 3, retaining 91 of 99 hybrids.

Although both controls yielded low error estimates, they are conceptually distinct and complementary: reversed-substrate decoys measure error in mapping *de novo* sequences to substrate decompositions, whereas same-mass entrapment measures the probability that an incorrect isobaric sequence survives competitive spectral filtering. Error control should therefore align with the specific failure mode under investigation: substrate-derived decoys for mapping errors in defined-substrate systems, and same-mass entrapment for spectrum-level false acceptance across both defined and open-proteome discovery regimes.

## Discussion

TransPepMS evaluates whether a reported non-canonical peptide sequence is superior to explicit same-mass (isobaric) alternatives under a specified evidence model. This framework distinguishes a candidate that merely *can* explain a spectrum from one uniquely supported by discriminating fragment ions. Calibration on simulated spectra established a three-ion margin at which no control peptide was falsely retained. Crucially, insufficient spectral evidence led predominantly to conservative abstentions ("unresolved") rather than false calls.

Distinguishing missing evidence from intrinsic sequence limits guides downstream follow-up. Where distinguishing ions are theoretically possible but absent, higher-depth reacquisition may resolve the candidate. However, where sequence pairs share near-identical backbones, such as the SHLV**EY**TPKT/ SHLV**YE**TPKT hybrid pair, which yields at most two distinguishing terminal ions, no amount of further b and y-ion sampling can overcome the three-ion threshold. These cases require orthogonal evidence (e.g., intensity predictions, synthetic standards, or internal ions). An "unresolved" call represents neither confirmation nor rejection.

Explicit sequence competition complements standard FDR approaches. Standard database searches evaluate candidates only within their search space, and target-decoy methods estimate dataset-level error without guaranteeing per-spectrum discrimination against unsearched isobars. Decoys need not reproduce the particular same-mass alternatives relevant to an individual noncanonical assignment. TransPepMS explicitly evaluates these targeted rivals and identifies the specific competitor driving a rejection. Furthermore, its error metrics target distinct failure modes: *same-mass entrapment* measures spectrum-level false acceptance, whereas *reversed-substrate decoys* measure sequence-to-protein mapping error ^16^. Neither should be interpreted as a universal error bound or automatically equated with the false-discovery rate among retained study candidates.

Unlike reference-competition tools such as PepQuery and PepQuery2 ^27, 28^, TransPepMS enumerates non-indexed isobaric alternatives, including reordered junctions and donor hybrids (Table 1). On the Liepe dataset ^5^, TransPepMS revealed that 33 of 200 PepQuery2-"confident" spliced calls were better explained by an isobaric canonical peptide (Supplementary Table ST10), not established as absent. In the insulin-spike data, the reordered and first-generation hybrid alternatives of VCGVCGYTPKT are not reference peptides and so lie outside reference competition entirely (Fig. 6); the reported sequence exceeded the first-generation competitor by the required margin, but its terminal-ion lead over the reordered hybrid remained below that threshold, and predicted intensity supplied secondary support. These examples preserve the distinction between primary discrimination and corroborating evidence.

**Table 1.**
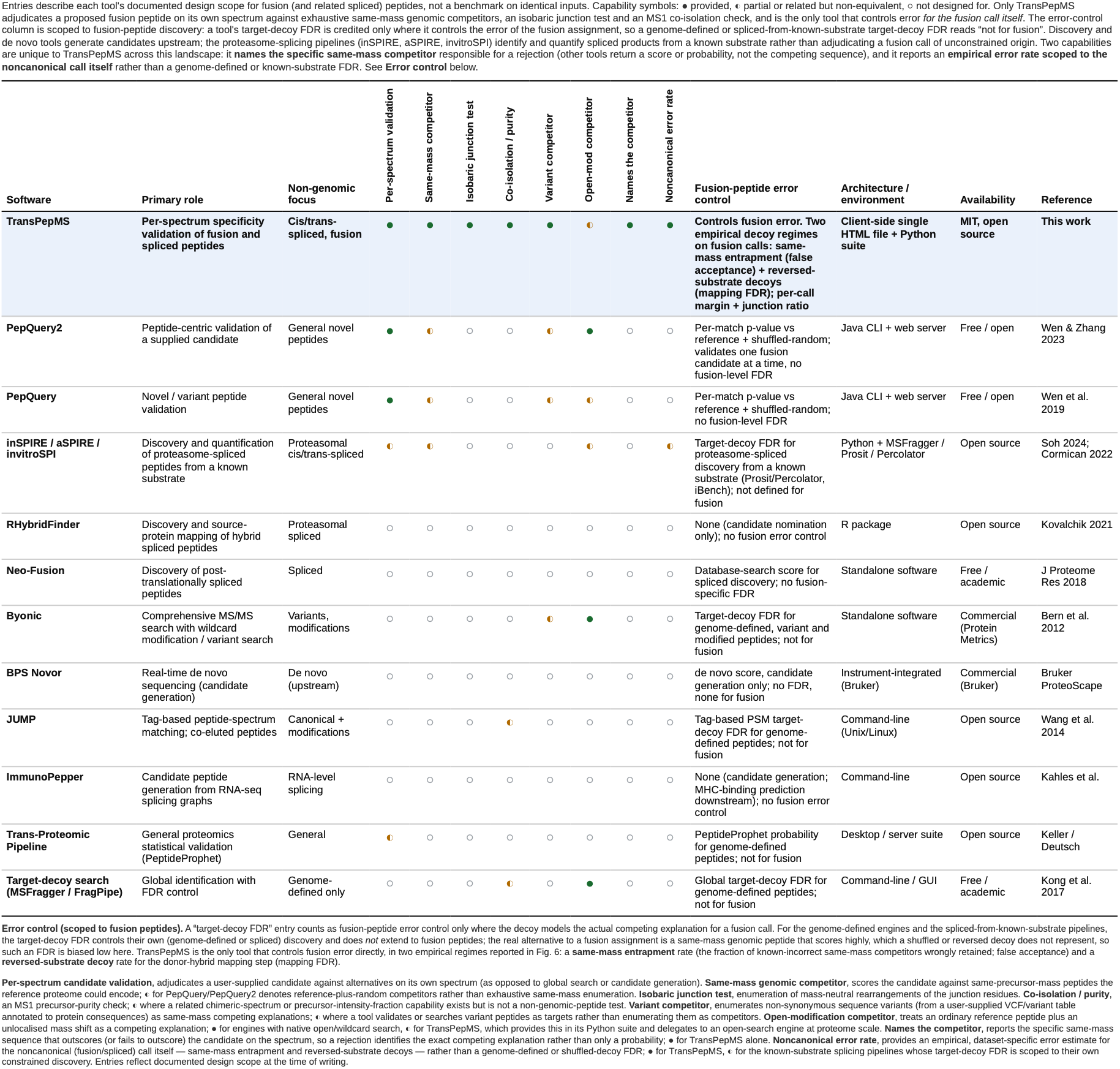
Overview of existing software scope, target-decoy handling, and error- control mechanisms for fusion-peptide identification compared with TransPepMS. Entries describe each tool’s documented design scope rather than performance on benchmarked inputs. System architecture, software availability, and error-model implementations for each tool are detailed in the respective columns of Table 1. Qualifiers and Methodological Differences. PepQuery and PepQuery2 perform competitive validation by submitting individual candidate peptide-spectrum matches, scoring them against reference proteomes and randomly shuffled decoys, and returning per-match confidence metrics. TransPepMS differs primarily in the structural composition of its competitor set, enumerating same-mass proteome peptides, junction rearrangements, and donor-constrained alternative hybrid peptides rather than randomized or shuffled decoys, and in identifying the specific competing sequence favoring a rejection; it represents an expanded, structurally constrained same-mass competition rather than a categorically distinct operation. The Trans-Proteomic Pipeline^42^ performs statistical validation of peptide-spectrum matches but does not construct explicit same-mass competitor pools. The JUMP software suite ^43^ resolves co-eluting precursor ions and chimeric mixture spectra, but does not implement precursor purity as a pre-filtering gate; similarly, while some target-decoy pipelines report precursor intensity fractions, these do not evaluate hybrid-peptide hypotheses directly. Byonic ^44^ searches reference databases using wildcard modification strategies rather than evaluating pre- specified candidate sequences. ImmunoPepper ^25^ generates candidate peptides from RNA-sequencing splice graphs, representing transcriptomic rather than proteasomal splicing, and relies on downstream MHC-binding predictions. Finally, standard target- decoy database searching cannot evaluate or score candidate sequences absent from the underlying reference database.

The applications establish different aspects of the tool’s scope. The fusion-neoantigen analysis separates candidate recovery, junction localization and same-mass competition; recovery alone does not establish either of the latter. In the contested spliced immunopeptidome, agreement with independent reanalyses ^17, 18^ shows that explicit competition reaches conclusions concordant with those reanalyses. This is concordance on shared spectra, not orthogonal experimental confirmation or evidence that rejected peptides are absent from the sample. Defined-donor reactions additionally permit competition against alternative hybrids. The deliberately mislabeled canonical controls show that independent database-search assignments and enumerated same-mass competitors can identify different incorrect calls.

The conclusions remain conditional on the set of candidates and the evidence model. A mass-unique assignment indicates that no competitor was found within the tested constraints, not that no alternative exists. The primary verdict uses terminal b/y-ion evidence; other measurements require separate validation before incorporation into that decision. The advisory precursor-purity check also leaves a distinction between sequence support within a scan and attribution of its fragments to a particular precursor. Calibration should be reassessed when fragmentation, mass accuracy, the range of modifications considered, or candidate class changes materially. The six-peptide fusion dataset demonstrates transfer to independently generated candidates but does not establish performance across fusion-neoantigen classes. Blinded evaluation using synthetic non-canonical peptides and matched same-mass competitors would provide a more stringent assessment.

TransPepMS makes the evidence underlying a non-canonical assignment inspectable: the distinguishing ions, tested alternatives, and reason for the reported verdict. This supports selection of candidates for further analytical and biological validation and identifies comparisons that require additional evidence. This framework accelerates candidate prioritization for downstream analytical verification, HLA-presentation assays, and functional T-cell validation. Establishing peptide production, HLA presentation and T-cell recognition remains a separate experimental task.

## Methods

### Sequence-Mapping Decoy False-Discovery Rate

For datasets generated from defined substrate systems, we estimated a class-specific false-discovery rate (FDR) for *de novo* cis/trans-spliced peptide identifications using a sequence-mapping target-decoy strategy ^16^. A *de novo* peptide, defined as having a PEAKS Average Local Confidence (ALC) score above the established threshold and at least a two-fold signal increase over control, was classified as cis/trans-spliced if it decomposed into two or more non-contiguous substrate fragments of at least four residues each. Leucine and isoleucine were treated as isobaric, and contiguous single-protein peptides were excluded.

The resulting *de novo* peptide pool was mapped simultaneously against target substrate sequences (mature human insulin and SARS-CoV-2 spike protein) and size-matched, sequence-reversed decoy substrates. The FDR was calculated as the ratio of peptides callable as spliced against the decoy set to those called against the target set. Decoy sequences must be strictly size-matched to the target proteins; a multi-proteome decoy is undefined for this framework because large database sizes naturally contain nearly every short k-mer by chance. This control requires a defined substrate set and is thus not applicable to open-proteome discovery, where spectrum-level same-mass entrapment was used instead.

### Spectral Hypothesis Competition Framework and Workflow

TransPepMS casts peptide identification as a competition among rival sequence hypotheses to explain an observed spectrum (Fig. 1 b). The pipeline evaluates candidate identifications (supplied by external search software) through three primary same-mass (isobaric) tests operating downstream of an advisory precursor-purity filter, alongside a secondary donor-hybrid test. Each test constructs *in silico* competitors of identical precursor mass to systematically rule out specific classes of alternative explanations:

1. **Test 1** (Independent Engine Competition): Competes the candidate sequence against the peptide assigned to the same scan by an independent database search (e.g., MSFragger/FragPipe psm.tsv or MaxQuant tables).
2. **Test 2** (Canonical Proteome Competition): Competes the candidate sequence against all same-mass peptides that reference proteomes could encode. This is run at two pool depths: unmodified canonical peptides, followed by a broadened pool containing common post-translational modifications (PTMs) or human missense variants.
3. **Test 3 (Junction Localization Test):** Competes the candidate sequence against all same-mass sequence rearrangements within a six-residue junction window.
4. **Test 4 (Exhaustive Donor-Hybrid Test):** Evaluates all alternative hybrid peptides that the specified donor proteins could form at the candidate precursor mass. For pairs tied on terminal-ion counts, predicted fragment intensity is applied as a secondary decision axis; intensity scoring can resolve comparisons within the unresolved margin but cannot override a primary three-ion rejection.

A candidate sequence is retained only if it maintains the required distinguishing-ion lead over competitors across all applicable primary tests (Tests 1-3).

### Precursor-Purity Filter

Before running primary tests, precursor purity is evaluated as the candidate isotope-envelope ion current divided by total ion current within the isolation window, extracted from centroided MS1 survey scans in mzML format (ProteoWizard MSConvert) ^35^. This metric corresponds to the precursor intensity fraction used in MaxQuant ^22^. Because quadrupole transmission and relative fragmentation efficiency are unmodeled, this MS1 window metric serves as an approximation of the precursor’s contribution to the MS2 spectrum. Precursor purity below 0.5 triggers an advisory flag for manual review but does not cause automated rejection.

### Candidate Evaluation Against Same-Mass Hypotheses

#### Test 1: Proteome and Database Competitor Assignment

Candidate sequences were derived from PEAKS Studio *de novo* identification ^20, 36^ or direct manual specification and verified as contiguous proteome substrings. When database search outputs (e.g., psm.tsv from MSFragger/FragPipe) from the same raw file were available, the corresponding assigned peptide was incorporated as a direct competitor. Reference database searches were conducted using FragPipe 22.0 (MSFragger 4.4.1, Philosopher 5.1.1 ^37^) against six target proteomes plus cRAP (199,720 total sequences). Search parameters included:

- Non-specific enzymatic digestion (up to two missed cleavages)
- Precursor tolerance: 15 ppm; Fragment tolerance: 20 ppm
- Selected ions: b and y-ion series
- Variable modifications: Methionine oxidation, N-terminal acetylation, glutamine pyroglutamate, and asparagine deamidation
- Decoys: Reversed sequences; Filter: 1% PSM false-discovery rate (FDR)
- Cysteine handling: Adjusted according to sample preparation (e.g., free cysteine for unalkylated insulin-Spike reactions)

#### Test 2: Exhaustive Same-Mass Enumeration and Rescoring

To identify competing sequences, Test 2 exhaustively enumerates all non-specific, same-mass peptides within the proteome databases under the modification set defined in Test 1. Each enumerated candidate is rescored against the candidate MS/MS spectrum using a modification-aware b and y-ion matcher. Localized modifications shift the theoretical masses of all fragment ions containing the modified residue (+15.9949 Da for methionine oxidation), allowing oxidized and unoxidized competitor ion series to be matched distinctly. Matches were evaluated up to the precursor charge state.

Mass tolerances were tailored to specific analyses:

- **Primary competition:** 10 ppm precursor and 15 ppm fragment (results remained stable between 10 and 15 ppm)

### • Modified and variant competitor broadening: 15 ppm

- **FragPipe database searching:** 20 ppm (fragment)
- **PepQuery2 searching:** 0.02 Da

Beyond the reference sequence pool, we expanded the search by re-enumerating all 109 target peptides to include asparagine/glutamine deamidation, methionine oxidation, and 79,334 curated human missense variants from UniProt *humsavar* ^38^. Variant peptides were restricted to those within 10 ppm of the candidate precursor mass and scored with modifications positioned to maximize the spectral match. None of the enumerated variant or modified competitors outscored any reported candidate peptide. On average, a median of 82,066 competitors were evaluated per spectrum (excluding the candidate’s own genomic entry).

Candidate validity was assessed using two sub-tests:

- **Test 2a (Total Matched-Fragment Comparison):** A zero-margin criterion evaluating whether any same-mass competitor matches the spectrum equally well or better based on total matched fragments. Results are reported as the fraction of competitors beaten or tied.
- **Test 2b (Distinguishing-Terminal-Ion Margin):** Evaluates the difference in unique diagnostic fragments between the candidate (kR) and its top same-mass competitor (kC). The best competitor is defined as the sequence yielding the smallest margin relative to the proposed candidate (ties are resolved by total matched-ion count).

Because complementary b and y-ions represent the same backbone cleavage event, counting is capped at two ions per unique fragmentation (2k). Thus, a reported sequence can lead an equal-length competitor by at most 2k distinguishing ions. We applied a fragmentation-resolved, peak-exclusive counting scheme in which each distinct fragmentation is counted once, and each observed peak is assigned to at most one ion. On the 109 insulin-Spike spectra, this corrected statistic agreed with raw matched-ion differences in 99 cases (differing by one ion in 10). On 4,374 iBench spectra, the corrected margin altered 423 comparisons (9.7%) and shifted the final verdict call in 39 spectra (0.9%). All reported margins use this corrected statistic. The canonical control on PXD000508 reports both: 3 of 200 spectra are ambiguous under Test 2a, and under Test 2b the known correct sequence leads its narrowest competitor by at least three distinguishing ions for 188 of 200 candidates.

The primary rule applies terminal-ion adjudication (b and y-ions only); internal and neutral-loss ions are considered secondary corroborating evidence. Test 1 uses an asymmetric decision rule: because database-assigned competitors represent identified genome-encoded peptides, candidate sequences are rejected if the competitor matches the spectrum equally well or better (rather than requiring a 3-ion lead).

#### Test 3: Junction Localization

Test 3 evaluates junction validity by enumerating all permutations within a 6-residue window surrounding the splice site (3 residues on either side). Where necessary, windows are expanded by mass-neutral residue swaps (e.g., N\GG). By evaluating all amino acid combinations up to length four by exact elemental formula, eleven elementary same-mass operations were defined alongside leucine/isoleucine equivalence (carbamidomethyl-cysteine vs. cysteine-glycine swaps applied only under fixed alkylation conditions). Junction confidence is quantified using the Junction Competitor Ratio (JCR):

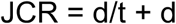

where d is the number of enumerated same-mass rearrangements scoring equal to or higher than the candidate, and t is the number of rearrangements outscored by the candidate. JCR = 0 indicates the candidate outscored all rearrangements. A junction is considered localized when JCR ≤ 0.05 and at least four rearrangements were enumerable (t + d **≥** 4).

### Decision Rules and Outcome Classifications

Outcomes are mutually exclusive, defined per candidate, and evaluate the fragmentation- resolved unique-ion margin (m) symmetrically:

- **Retained (m ≥ +3):** Requires m **≥** +3 (and no competitor leading on union spectral angle beyond the abstention band when intensity scoring is applied). A margin threshold of +3 was selected via simulation to exceed the maximum possible terminal-ion margin of a single-fragmentation competitor (2k = 2).
- **Unresolved (-2** ≤ **m** ≤ **+2):** Includes exact ties (m = 0). Unresolved spectra are categorized using specific reason codes in the Supplementary Data (sheet S3):
- *Structurally non-identifiable pair* (no diagnostic terminal ions exist)
- *Insufficient diagnostic fragmentation* (diagnostic ions absent or undetected)
- *Spectral tie* (identical matched-ion count)
- *Intensity-axis abstention* (within the decision band)
- *Low-purity flag*
- **Rejected (m** ≤ **-3):** Candidates outscored by a competitor by 3 or more distinguishing ions.
- **Mass-Unique:** Candidates with no qualifying same-mass competitors in the database.

Co-isolation flags and synthetic-standard spectrum matches serve as advisory metadata and do not independently determine acceptance or rejection. Full sweeps across thresholds from 1 to 8 distinguishing ions are reported for completeness.

#### Test 4: Exhaustive Enumeration of Alternative Hybrid Peptides

Test 4 competes a proposed hybrid (fusion) peptide against the full space of alternative hybrid peptides that the same donor proteins could generate at the same precursor mass. This is the space Tests 1 to 3 cannot reach: the nearest competitor to a hybrid peptide is not a genome-encoded peptide but another hybrid of the same two donors, and such sequences are absent from every reference proteome and from the junction- rearrangement set of Test 3. The donor proteins are specified per dataset (for the insulin- Spike cohort, mature human insulin and the SARS-CoV-2 Spike glycoprotein; for a reported fusion neoantigen, its two fusion-partner proteins).

For each spectrum, the competitor library was built directly from the donor sequences in two steps. First, every contiguous substring of 2 to 15 residues was extracted from each donor as a candidate fragment, representing any stretch a donor could contribute to a hybrid. Second, these fragments were concatenated, up to three at a time, in any order and from either donor, to assemble candidate hybrid products. Joining two fragments creates a single junction (a first-generation hybrid) and joining three creates two junctions (a second-generation hybrid); fragments taken from the same protein give *cis* configurations and fragments from different proteins give *trans* configurations, so all *cis*, *trans*, first- and second-generation arrangements are represented. Only products of 8 to 25 residues whose theoretical precursor mass fell within 10 ppm of the measured precursor were kept as same-mass competitors for that spectrum.

The enumeration is exhaustive within this donor-constrained construction: every sequence the named donors can build under the stated length and mass limits is generated and scored, rather than a sampled subset, although it is not every conceivable peptide of that mass. Because the number of qualifying three-fragment arrangements grows combinatorially with donor length, the competitor set reaches into billions of configurations per spectrum for the larger donors. Test 4 is therefore implemented in a compiled scoring kernel and parallelized across high-performance-computing nodes, with a pure-Python reference implementation retained for verification.

Each competitor, and the reported peptide, were scored against the same experimental tandem mass spectrum using the modification-aware b and y-ion matcher of Tests 1 and 2, at 15 ppm fragment tolerance and over charge states from 1 to the precursor charge. The distinguishing-ion count was the fragmentation-resolved, peak-exclusive count defined for Test 2b, so each backbone cleavage contributes at most two ions (a b-ion and its complementary y-ion) and each observed peak is assigned to at most one ion. For every spectrum, Test 4 records the reported-peptide score, the best competitor and its score, the distinguishing-ion margin between them, the total number of enumerated competitors, the count of competitors matching or exceeding the reported score, and the full competitor-score histogram.

The best alternative hybrid is almost always a single-residue re-spelling of the reported sequence, an adjacent transposition or a mass-equivalent block substitution, differing from it at only one backbone cleavage. At most two distinguishing ions can therefore separate the two sequences, so the margin at Test 4 cannot reach the three-ion retention threshold, and Test 4 does not by itself retain or reject a hybrid on terminal-ion count. For the pairs that terminal-ion counting leaves tied or separated by fewer than three ions, predicted fragment intensity (the spectral-angle axis described in the next section) is applied as a secondary decision axis; it can resolve comparisons within the unresolved band but never overrides a primary three-ion rejection from Tests 1 to 3.

Because products may combine up to three fragments, a reported second-generation (two-junction) assignment is competed directly against its first-generation isobars and is retained only when the fragment data explicitly localize both junction sites. After rescoring and leucine/isoleucine consolidation, each top-ranking competitor was classified as a sequence reordering, an isobaric block substitution, or a compositionally distinct sequence.

### Secondary Evidence from Predicted Fragment Intensity

Predicted fragment ion intensities served both as per-spectrum corroboration for retained identifications and as an orthogonal decision axis for candidate pairs unresolved by matched ion counts. All predictions were generated using CPU implementations of the pre-trained AlphaPeptDeep models (v1.5.0) ^39^. Model performance on non-tryptic peptides was independently validated using a held-out dataset (Supplementary Fig. S2). Because instrument collision energy settings vary across acquisition runs, nominal energies were empirically calibrated by scoring high-confidence spectra against their known sequences across an energy grid. The setting yielding the maximum median self- spectral angle was selected for downstream analysis (30 for the insulin-Spike dataset and 35 for the calibration dataset).

Candidate sequences were compared using the spectral angle (SA) evaluated across the union of their predicted fragment positions: SA = 1 − (2/π) arccos (p^ · ô) where p^ and ô are the L2-normalized predicted and observed intensity vectors, respectively. Evaluating the union of predicted positions, rather than restricting the calculation to each candidate’s specific positions, prevents isobaric substitutions from receiving artificially inflated scores due to unassigned positions.

To establish error rates prior to scoring experimental hybrid-peptide spectra, decision thresholds were calibrated on two independent ground-truth control sets:

1. **HeLa dataset:** 1,388 peptides from PXD000508 ^33^
2. **Synthetic peptide pool:** Chemically synthesized peptides from ProteomeTools (PXD021013) ^40^

For each reference peptide, decoy variants were generated via adjacent residue transpositions and isobaric block substitutions. Decoys were scored blindly against the target sequence, and null distributions were constructed by scoring pairs of decoy variants against one another.

The decision threshold was fixed exclusively on the subset of spectra where candidate ion counts resulted in a tie or a loss, and was subsequently applied without modification:

- **HeLa calibration set:** At a threshold of 0.08, the rule resolved 43.6% of tied cases with a 1.0% error rate (11 of 1,101 calls; 95% bootstrap CI: 0.5%-1.6%, seed

20260726).

- **ProteomeTools synthetic set:** At the same threshold, the rule achieved a 48.0% decision rate with a 0.7% error rate (17 of 2,421 calls; 95% bootstrap CI: 0.3%-1.2%).

The AlphaPeptDeep model was evaluated strictly as released, without fine-tuning on any hybrid-peptide spectra. All decision thresholds were locked prior to evaluating experimental insulin-Spike data. Because comparisons sharing a common spectrum or sequence violate independence assumptions, proportion confidence intervals were calculated using non-parametric bootstrap resampling blocked by whole peptides; one-sided upper confidence bounds for zero-event or rare-event rates, for which a resampling interval is degenerate, were obtained from the exact binomial (Clopper-Pearson) distribution.

### Manual Inspection and Orthogonal Validation

Peptide-spectrum matches (PSMs) retained after automated pairwise competition were subjected to expert manual inspection using a dedicated interactive visualization tool. This tool mapped theoretical junction-spanning internal fragments against observed peaks to confirm covalent continuity across splice sites and rule out alternative structural rearrangements. Primary diagnostic identifications were based strictly on terminal b and y-ions, whereas internal fragments provided corroborating evidence for backbone connectivity.

When synthetic peptide standards were available, experimental spectra from enzymatic digests were validated against synthetic reference spectra by calculating cosine similarity over matched fragment intensities using a pre-specified low-mass cut-off. Spectral match scores and retention-time alignments are reported individually per candidate peptide as independent supporting evidence and were kept separate from the primary competition score.

### In Silico Performance Evaluation Using a Simulated Benchmark

To systematically evaluate the performance of the pairwise same-mass competition rule under controlled parameters, we generated a synthetic benchmark dataset using a dedicated simulation framework (TransPepMSsim.py, deterministic seed 20260805). Synthetic datasets with known ground truth provide an established benchmark for evaluating peptide identification algorithms ^31^.

#### Experimental Design and Factorial Stratification

The benchmark used a fully crossed, six-factor design yielding 864 unique strata, with 1,200 simulated cases per stratum (1,036,800 total spectra):

1. **Peptide length:** 9, 11, 13, or 15 residues
2. **Precursor charge state:** +2 or +3
3. **Fragment recovery rate:** 0.50, 0.70, or 0.90 (proxy for collision energy efficiency)
4. **Co-isolation contamination:** 0%, 20%, or 50% contaminant fragment ion current
5. **Competitor structural class:** Six distinct competitor types:

1. Single adjacent residue transposition
2. Double adjacent residue transposition
3. Triple adjacent residue transposition
4. Mass-neutral isobaric block substitution
5. Same-composition structural anagram
6. Unrelated peptide selected from a mass-indexed database of 400,000 random sequences within 10 ppm of the target precursor mass
6. **Generating sequence identity:** Target sequence or same-mass competitor sequence Leucine and isoleucine substitutions were excluded from all competitor classes.

A stratum is one unique combination of the six factor settings, formed by choosing one level from each factor; for example, a 9-residue peptide at +2 charge, 0.50 recovery, 0% co-isolation, a single adjacent-transposition competitor, and target generating identity is one stratum, and changing any single setting defines a different stratum. Within each stratum, 1,200 spectra were simulated that share that condition but differ only by random noise, such as which fragment ions are recovered and where contaminant signal falls. The six factors are fully crossed, so the number of strata is the product of their levels: 4 lengths × 2 charge states × 3 recovery rates × 3 co-isolation levels × 6 competitor classes × 2 generating identities = 864 strata. With 1,200 simulated cases in each stratum, the benchmark contains 864 × 1,200 = 1,036,800 spectra in total. The sixth factor, generating sequence identity, splits these spectra into two arms of equal size that are analyzed separately. In the correct-candidate arm the spectrum is generated from the target sequence, so the target is the correct answer and the quantity of interest is how often it is retained. In the incorrect-candidate arm the spectrum is generated from the same-mass competitor, so the target is a known-incorrect answer and the quantity of interest is how often it is nonetheless retained, that is, falsely accepted. Holding the generating identity fixed removes that one factor and leaves the other five, 4 × 2 × 3 × 3 × 6 = 432 strata; at 1,200 cases each, every arm therefore contains 432 × 1,200 = 518,400 spectra, exactly half of the 1,036,800 total. Each arm is scored independently, and every per-margin statistic reported below is computed over the 518,400 spectra of a single arm.

### Spectral Generation and Automated Scoring

Spectra were synthesized using an independent fragmentation model decoupled from the scoring algorithm. The generator adjusted individual fragment ion yields based on local sequence context, emitting singly charged ions universally and doubly charged ions with a probability proportional to the precursor charge state. Injected spectral variability included:

- Per-spectrum mass offsets sampled from N (0, 4 ppm)
- Per-peak measurement jitter sampled from {N (0, 3 ppm)
- Uniform background noise
- Co-isolated contaminant fragments (where parameterized)

The automated scorer evaluated peak lists against target and competitor sequences using b and y-ions only, applying a 15 ppm match tolerance. Distinguishing fragment positions were explicitly calculated for each pair, counted once across charge states, and restricted to a single peak assignment per ion.

#### Error Analysis and Benchmark Performance

Because control sequence identities were known for all simulated spectra, candidate retentions were categorized as correct or false assignments to determine error rates across ion-margin thresholds. Error rates are reported with 95% Wilson score confidence intervals (zero-numerator cases are reported as a one-sided 95% upper bound of 3/N). Across all 1,036,800 simulated test cases, the unique-ion margin separated target and competitor distributions with an overall receiver operating characteristic area under the curve (ROC-AUC) of 0.949:

- **1-ion margin:** False-retention rate of 3.6 x 10^-3^
- **2-ion margin:** False-retention rate of 5.2 x 10^-5^
- **≥3-ion margin:** Zero false retentions observed across all 518,400 evaluated cases in the incorrect-candidate arm Detailed performance metrics for all individual strata and thresholds are provided in the Supplementary Data workbook. This synthetic benchmark characterizes the mathematical behavior of the same-mass rule under explicit modeling assumptions; real-world empirical validation on acquired experimental spectra was performed using the iBench ground-truth dataset.

### Empirical Validation Datasets and Micro-Environment Benchmarks

#### Benchmark Datasets (iBench)

Empirical validation of the same-mass competition framework was performed using the iBench control benchmark dataset (PXD044451) ^32^. This dataset comprises 4,374 spectra acquired on an Orbitrap Fusion Lumos mass spectrometer from 24 h digestions of five synthetic polypeptide substrates by the human 20S proteasome, yielding 1,183 unique peptide sequences across 2,519 canonical, 1,398 *cis*-spliced, and 457 *trans*-spliced peptide-spectrum matches (PSMs).

Non-specific competitor sequences were exhaustively enumerated across six reference human proteomes plus cRAP using a precursor mass tolerance of 10 ppm and a fragment mass tolerance of 20 ppm, yielding a median of 2.0 x 10^5^ candidate sequences per spectrum. Performance was evaluated across two analytical units (specified in corresponding figure captions):

- **Per-spectrum analysis (kK binning):** Evaluated over the 1,911 spectra possessing at least one enumerable same-mass competitor (Fig. 2d).
- **Per-sequence analysis (length and substrate stratification):** Evaluated across the 1,183 unique peptide sequences by taking the median per-spectrum result for each sequence (Supplementary Fig. S8).

#### Cumulative-Mass Fragment Counting and Distinction Metrics

To maintain rigorous definitions when comparing sequences of differing length, distinguishing-ion counts were defined on cumulative backbone fragment masses rather than linear amino acid residue positions. Each candidate peptide is represented by its set of internal backbone cleavage masses (with leucine and isoleucine treated as equivalent), matched within a tolerance of 0.02 Da.

For a target sequence R and its competitor C, kR and kC denote the number of cumulative fragment masses unique to R and C respectively (where R is a known benchmark sequence, kK is substituted for kR):

- **Equal-length competitors:** kR = kC, both equaling the number of backbone positions where the cumulative-mass series differ.
- **Unequal-length competitors:** The metric is inherently asymmetric. For example, replacing an asparagine (N) with a glycine-glycine (GG) dipeptide preserves the precursor mass; however, GG contains an internal cleavage site between the two glycine residues that N lacks. Thus, kR = 0 and kC = 1.

The deployed unique-ion margin incorporates this structural asymmetry by subtracting unique competitor fragment ions from unique target sequence fragment ions. Consequently, a candidate peptide lacking unique fragment masses (kR = 0) can at best achieve a tie, but can never lead a competitor. Because peptides derived from the same synthetic substrate do not represent independent observations, consistency of effect across the five substrate groups was confirmed using a one-sided exact sign test on paired substrate observations.

### Sequence-Imposed Discrimination Limits

Consider a candidate sequence R and a precursor-isobaric competitor C evaluated under a fixed fragmentation model (b and y-terminal ions, specified fragment tolerance, and Leu/Ile equivalence). Let kR and kC represent the number of unique cumulative backbone fragment masses for R and C, respectively.

Because precursor mass is fixed, each unique fragment mass yields at most one b-ion and one complementary y-ion. Consequently, sequence R can contribute at most 2kR distinguishing terminal ions, while competitor C can contribute at most 2kC. The peak- exclusive distinguishing-ion margin (m) for any spectrum derived from this precursor is strictly bounded:

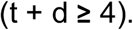

These theoretical bounds depend solely on sequence composition and the theoretical model, not on acquired spectral quality. A definitive verdict is attainable only when the theoretical upper bound meets or exceeds the decision threshold (m**≥**3):

- Retaining candidate R requires 2kR **≥** 3 (kR **≥**2).
- Rejecting candidate R in favor of competitor C requires 2kC **≥**3 (kC **≥** 2). Candidate pairs where both kR ≤1 and kC ≤1 and are structurally unresolvable by terminal ions under this model, regardless of spectral signal-to-noise ratio, dynamic range, or fragmentation coverage.

### Experimental Hybrid-Peptide Validation and Control Datasets

#### Experimental Hybrid-Peptide Dataset

Validation data for hybrid peptides were obtained from a companion study of cathepsin-mediated reverse proteolysis ^16^, comprising 117 PSMs from unalkylated reactions between mature human insulin and SARS-CoV-2 Spike glycoprotein. These represent 109 unique peptides, each evaluated against its corresponding experimental spectrum. When a first-generation sequence was identified across multiple PSMs, the single spectrum carrying the highest PEAKS Average Local Confidence (ALC) score was selected prior to same-mass competition to prevent selection bias. Distinguishing-ion margins were calculated independently per spectrum across all 109 cases. For sequence-level summaries, these collapse to 105 distinct sequences under Ile/Leu equivalence, aggregated using the median margin across spectra. *De novo* candidates exported from PEAKS Studio were filtered at an ALC threshold of **≥** 90%.

#### Positive, Negative, and Calibration Controls

- **Resolution-Matched Negative Controls:** As a negative control, 46 canonical PSMs were deliberately relabeled as hybrid peptides and processed through the identical analytical cascade alongside true hybrid candidates, forming the 145 evaluation cases in Fig. 5 a. Planted control peptides were classified as correctly rejected when either a verbatim proteome entry (Test 1) or a superior genome-encoded competitor (Test 2) accounted for the spectrum.
- **Predicted-Intensity Calibration Sets:** Predicted fragment intensity models were calibrated using two independent datasets of genome-encoded peptides containing no spliced or fusion junctions: HeLa dataset PXD000508 ^33^ (serving also as a negative control) and the ProteomeTools synthetic peptide pool (PXD021013) ^40^.

#### Independent Re-Evaluation with PepQuery2

Independent spectrum-level validation was performed using PepQuery2 (v2.0.2) in PSM- validation mode ^27, 28^. Each candidate sequence was submitted alongside its corresponding experimental spectrum as a targeted query against the six-proteome reference database supplemented with cRAP contaminants. Search parameters were configured as follows:

- Enzymatic specificity: Tryptic digest
- Precursor mass tolerance: 10 ppm
- Fragment mass tolerance: 0.02 Da
- Precursor charge states: Up to +2
- Variable modifications: Methionine oxidation
- Fixed modifications: Dataset-matched cysteine alkylation status

For the head-to-head comparison (**Fig. 6**), every candidate spliced peptide-spectrum match (PSM) was evaluated concurrently using both PepQuery2 and the TransPepMS cascade on identical raw spectra. A PepQuery2 assignment was classified as confident if retained after competition against same-mass reference peptides and a random-sequence null distribution (utilizing PepQuery2’s default significance threshold).

PepQuery2 yielded 200 confident assignments across the evaluation set. On these identical 200 spectra, the TransPepMS distinguishing-ion margin (m), applying decision thresholds of retained (m **≥** +3), unresolved (-2 ≤ m ≤ +2), and rejected (m ≤ -3), retained 33 candidates, classified 134 as unresolved, and rejected 33 candidates. Each of the 33 rejections identified a specific genome-encoded, same-mass competitor sequence that explained a greater number of distinguishing fragment ions than the proposed spliced sequence (e.g., scan 30840, where the reported spliced peptide GPPGSVILY was outscored by the genome-encoded sequence ILGPPGSVY; Fig. 6b). Comprehensive per-spectrum ion assignment tables for all 200 head-to-head comparisons are provided in the reproducibility repository.

#### External Validation on the GF-NEO Immunopeptidome Dataset

External applicability was evaluated using the GF-NEO dataset (PXD063253), which investigates pediatric cancer fusion-junction hybrid peptides resulting from chromosomal translocations [26]. Six reported junction peptides corresponding to ETV6-RUNX1 and CBFB-MYH11 fusion events were evaluated against deposited peak lists using fixed carbamido-methylation on cysteine and variable oxidation on methionine. Peptides were queried using a precursor mass tolerance of 15 ppm and a fragment mass tolerance of 20 ppm (b and y-ions).

All six reported fusion peptides were successfully detected in the peak lists. The five peptides exhibiting the highest sequence coverage were advanced to same-mass competition (Test 2) and junction localization (Test 3); candidate MPIGRIADA (7 matched fragment ions) was excluded due to insufficient coverage. Each advanced sequence was evaluated under Test 3 with the junction position defined by the reported genomic fusion breakpoint.

Because the repository supplies processed peak lists rather than raw instrument files or survey scans:

1. **Test 1** (independent database search) and MS1 co-isolation filtering could not be performed.
2. **Test 4** (donor fragment recombination) is conceptually inapplicable to chromosomal translocation junctions, where the primary competing hypothesis is an alternative breakpoint rearrangement (evaluated in Test 3).
3. **Fragment intensity rescoring** was omitted, as class-matched intensity calibration models for non-tryptic Class I immunopeptidomes are not currently available.

For Test 2, same-mass competitors were exhaustively enumerated across the six reference human proteomes plus cRAP, incorporating fixed cysteine carbamido- methylation and up to two variable methionine oxidations. Enumerated competitor pools ranged from 17,210 to 30,999 sequences per spectrum (median 17,210).

### Re-Adjudication of a Published Hybrid-Peptide Immunopeptidome

To re-evaluate previously published hybrid-peptide identifications, we retrieved the 1,214 hybrid peptide-spectrum matches (PSMs) reported for dataset PXD000394 ^5^, alongside 3,603 non-spliced canonical identifications from the same raw files used as calibration controls, referencing scan numbers established in a subsequent reanalysis ^19^.

The raw instrument file was converted using ThermoRawFileParser (v1.4.0) ^41^ into peak lists for same-mass competition (Test 2) and junction localization (Test 3), and into mzML format retaining MS1 survey scans to evaluate the precursor co-isolation gate. Scan numbering was validated against reported precursor masses. Search parameters were set as follows:

- **Cysteine alkylation:** Unalkylated (free cysteine)
- Precursor mass tolerance: 10 ppm
- **Fragment mass tolerance:** 20 ppm
- **Peptide length filter:** 8-12 residues (HLA Class I immunopeptidome window)
- **Variable modifications:** 0-1 methionine oxidation
- **Database scope:** Six human reference proteomes plus cRAP contaminants Junction indices for Test 3 were defined directly from the splice coordinates provided in the original study. The co-isolation intensity threshold was set to 0.5 (the default advisory threshold). The co-isolation gate serves as advisory metadata and did not alter any sequence calls.

The predicted fragment intensity axis was excluded for this dataset due to high empirical error rates on peptides of established identity (16.4%-22.0%, compared to 1.0% on the HeLa control dataset). Decision calls from five independent reanalyses (four search engines: PEAKS, SPIDER, MaxQuant and Comet) were benchmarked per spectrum. Full analysis scripts and workflows are available in repository directory 13_spliced_reanalysis.

### Entrapment False-Acceptance Rate Estimation

To quantify the rate at which the analytical cascade incorrectly retains false hybrid-peptide assignments, known-incorrect candidate sequences were evaluated using the identical filtering gates and tests applied to experimental targets. For each high-confidence canonical PSM (defined as a genome-encoded peptide accepted at a 1% false discovery rate by an independent database search), a decoy candidate of identical precursor mass was constructed by joining two reference-proteome fragment sequences whose combined mass matched the canonical precursor within dataset tolerances. Decoy products occurring verbatim in the reference proteome or matching the true canonical sequence were excluded.

Each synthetic decoy candidate was assigned to the corresponding experimental spectrum and evaluated using the distinguishing-ion margin (m). Because the true canonical peptide is included in the enumerated same-mass competitor pool, a decoy candidate was retained only if its matched b and y-fragment ion count exceeded all same-mass alternatives, including the true sequence, by at least 3 distinguishing ions (m **≥**+3). Thus, false retention represents a spectrum-level misassignment rather than a sequence-level artifact. The false-acceptance rate (FAR) is defined as:

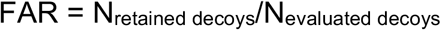

and is reported with 95% Wilson and one-sided Clopper-Pearson confidence intervals. Canonical peptides analyzed within the same instrument run served as paired positive controls.

Tests 1 through 3 were applied without co-isolation or predicted-intensity filtering; because those additional gates can only reduce candidate retention, the reported false-acceptance rates represent conservative upper bounds. For the PXD000394 immunopeptidome (Fib-2 run), 3,592 same-mass decoy candidates were generated from confident canonical PSMs against the reviewed human proteome and evaluated using exhaustive competitor pools (6 ppm precursor tolerance, 20 ppm fragment tolerance, free cysteine). The 3,603 canonical PSMs from the same run served as the paired positive control. Detailed candidate counts, confidence intervals, failure modes, and the single retained decoy case are provided in Supplementary Note S1.

## Statistical Analysis and Confidence Bounds

For Test 2, competitor overlap is expressed as the fraction of enumerated same-mass competitors matching or outscoring the candidate sequence per peptide. For zero-count outcomes, upper confidence bounds are reported using a one-sided Clopper-Pearson 95% interval.

Error rates for predicted fragment intensity thresholds are reported as proportion of incorrect calls among all decision-eligible spectra, with confidence bounds for non-zero rates determined via 2,000 bootstrap replicates resampled by whole peptide units to account for intra-sequence variance, and one-sided upper bounds for zero-event rates taken from the exact binomial (Clopper-Pearson) distribution. All statistical execution code, analysis scripts, and source data are archived in the public repository release.

## Implementation and Software Availability

TransPepMS is implemented as a lightweight, single-file HTML application that operates locally within any modern web browser without requiring a backend server or external network calls; all input data remain securely held in local browser memory. For high-throughput batch processing, TransPepMS is additionally distributed as an automated, cluster-scalable pipeline packaged within an Apptainer container alongside SLURM submission scripts. This headless workflow accepts user-supplied spectral files (.mzML or .mgf), candidate sequence tables, and reference proteome FASTA files, returning per-peptide decision verdicts and comprehensive summary reports without browser intervention.

Execution requires three core inputs: spectral data provided as .mzML or .mgf files, a candidate table specifying target peptide sequences and their corresponding scan indices, and one or more target proteome FASTA files. Optional candidate table columns allow users to specify precursor charge states, previously assigned database peptides for Test 1, and junction coordinates for Test 3, while supplying matching indexed .mzML files enables the precursor-purity gate by reading isolation window parameters directly from MS1 survey scans. Candidate sequence modifications written in bracketed notation are parsed to their bare amino acid backbones, with modification masses assigned to precursor and competitor pool masses rather than individual fragment matches; enumerated same-mass competitors bearing localized modifications are instead rescored using modification-shifted fragment ion masses under Test 2.

The Apptainer container fixes the computational environment, ensuring full reproducibility across HPC clusters without local installation dependencies while cluster scheduling is managed automatically via SLURM scripts. Cluster-scale parallel processing is deployed for Test 4, as its exhaustive hybrid-peptide competition evaluates up to billions of alternative configurations per spectrum. The core same-mass scoring algorithm relies on a compiled C-kernel for optimal performance, accompanied by a pure Python reference implementation that yields identical results when C-compilation is unavailable; both implementations and overall decision logic are validated through automated unit tests. An accompanying Python analysis suite evaluates spectral alignment between observed spectra and AlphaPeptDeep intensity predictions for retained candidates. Pipeline execution generates a structured tabular report detailing one row per candidate: the final verdict, top competitor sequence, and distinguishing-ion margin (m). Version 1.00 is open-source and archived under the MIT license with a version-tagged release, and the repository DOI (Zenodo) will be finalized prior to publication. Per-peptide cascade results, PepQuery2 runs, benchmark evaluations, negative controls, and secondary analyses are provided in the Supplementary Data Workbook (Sheets S1-S21).

## Data Availability

All mass spectrometry raw files and search outputs have been deposited to the ProteomeXchange Consortium via PRIDE:

- Insulin-Spike reverse-proteolysis dataset: PXD075848
- **Negative control and intensity calibration datasets:** PXD000508 and PXD021013
- **GF-NEO immunopeptidome dataset:** PXD063253
- **Re-adjudicated immunopeptidome datasets:** PXD000394 and PXD009660 Reference proteome sequences and human missense variants were retrieved from UniProt (*humsavar*). Detailed dataset descriptions, analytical parameters, and provenance metadata are provided in Supplementary Methods. Per-peptide cascade results, benchmark performance evaluations, control analyses, and intensity rescoring outputs are available in the Supplementary Data Workbook (Sheets S1-S21). Repository accession numbers and the Zenodo DOI will be finalized prior to publication.

## Code Availability

The interactive TransPepMS application, core cascade scorer, same-mass enumerator, variant parser, and intensity calibration pipelines are open-source and archived under the MIT license. The full repository, including high-performance C-kernels, SLURM cluster submission scripts, self-test suites, and figure generation code, is version-tagged, verified with SHA-256 checksums, and will be deposited on Zenodo upon acceptance.

Exhaustive hybrid-peptide competition workflows and intensity-discrimination pipelines are organized in dedicated directories (10_test5_exhaustive and 11_intensity_discrimination, respectively), each containing a README file detailing software dependencies, execution order, inputs, and limitations. Source code, release tags, and Zenodo DOI links will be finalized upon manuscript acceptance.

## Competing Interests

The authors have filed a provisional patent application relating to the method described in this manuscript. The authors declare no other competing interests.

## Use of Large Language Models

AI tools (Anthropic Claude, OpenAI GPT) were used for English-language editing and code debugging. The authors reviewed all output and take full responsibility for the contents of this manuscript.

## Notes

### Summary of Updates

This version corrects the placement of one Results paragraph. A paragraph describing the modified-competitor screen and the head-to-head comparison with PepQuery had been duplicated into the subsection that validates the three-ion rule on independent reference spectra, where it did not belong. That paragraph has been removed from that subsection, and the single summary statistic it contained has been moved to the Direct Comparison with PepQuery section, where it fits. No results, numbers, figures, or datasets were changed by this correction. Minor editorial and formatting corrections were also made for clarity and consistency.

### Funding

- Canadian Institutes of Health Research grants: PJT-155979

