## Supplementary material for "TransPepMS: A Tool to Verify Peptides Absent from Reference Proteomes": Supplimentary Figures 1-9

Spectral hypothesis competition tests identifications of peptides absent from reference proteomes

### Supplementary Figures

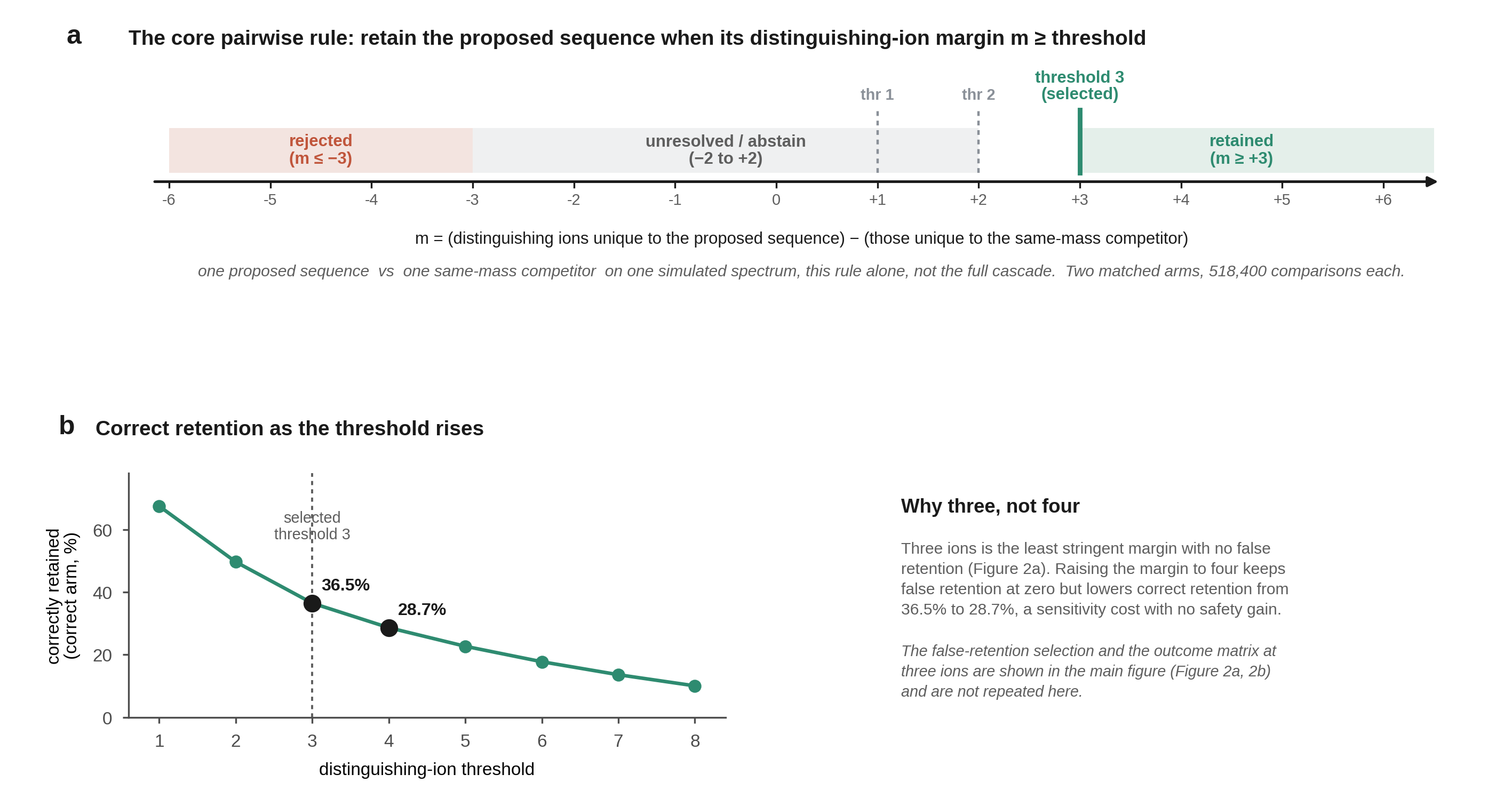

**Supplementary Figure S1. The core pairwise decision rule and the sensitivity cost of the threshold (1,036,800 simulated cases).** Every spectrum is generated by a simulator; no acquired spectrum enters this figure, and the figure characterizes the pairwise rule alone, not the complete cascade. The threshold selection by false retention and the outcome matrix at three ions are in the main figure (Figure 2a and 2b) and are not repeated here. Two matched arms of 518,400 comparisons each: a correct-candidate arm (the true sequence submitted) and an incorrect-candidate arm (a wrong sequence submitted). (a) The decision rule on the distinguishing-ion margin m, the ions unique to the proposed sequence minus those unique to the same-mass competitor, with candidate thresholds of one, two, and three ions. (b) Correctly retained comparisons in the correct arm against the threshold, 36.5% at three ions and 28.7% at four, so raising the margin to four ions lowers sensitivity without removing any false retention, which is why three is selected.

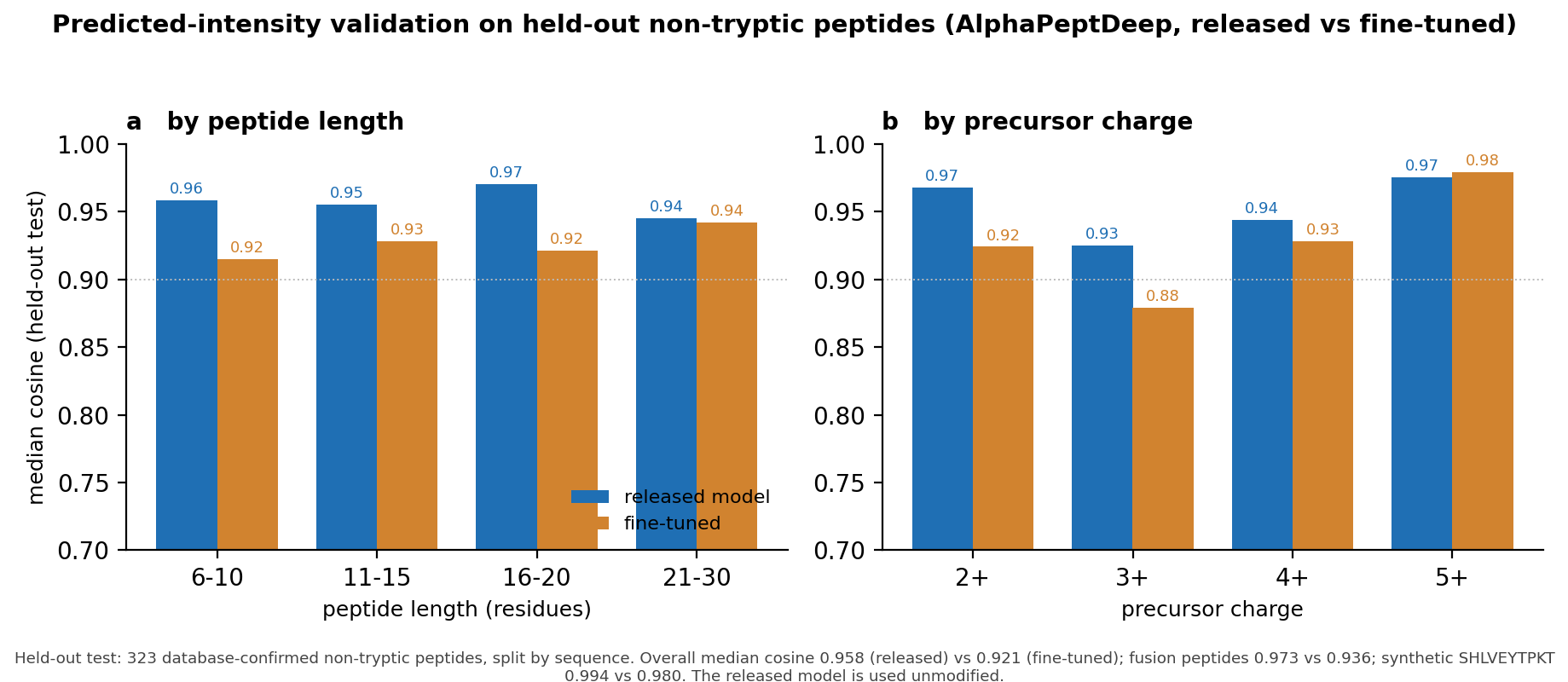

**Supplementary Figure S2.** Predicted-intensity validation on held-out non-tryptic peptides. Overall median cosine 0.958 (released) against 0.921 (fine-tuned) on 323 held-out peptides, and 0.973 against 0.936 on the fusion-junction hybrid peptides; across the three synthetic standards the released model reached 0.994 for SHLVEYTPKT, 0.976 for LVCGFVNQHLC and 0.843 for AENSVAYTPKT. The released model is used unmodified.

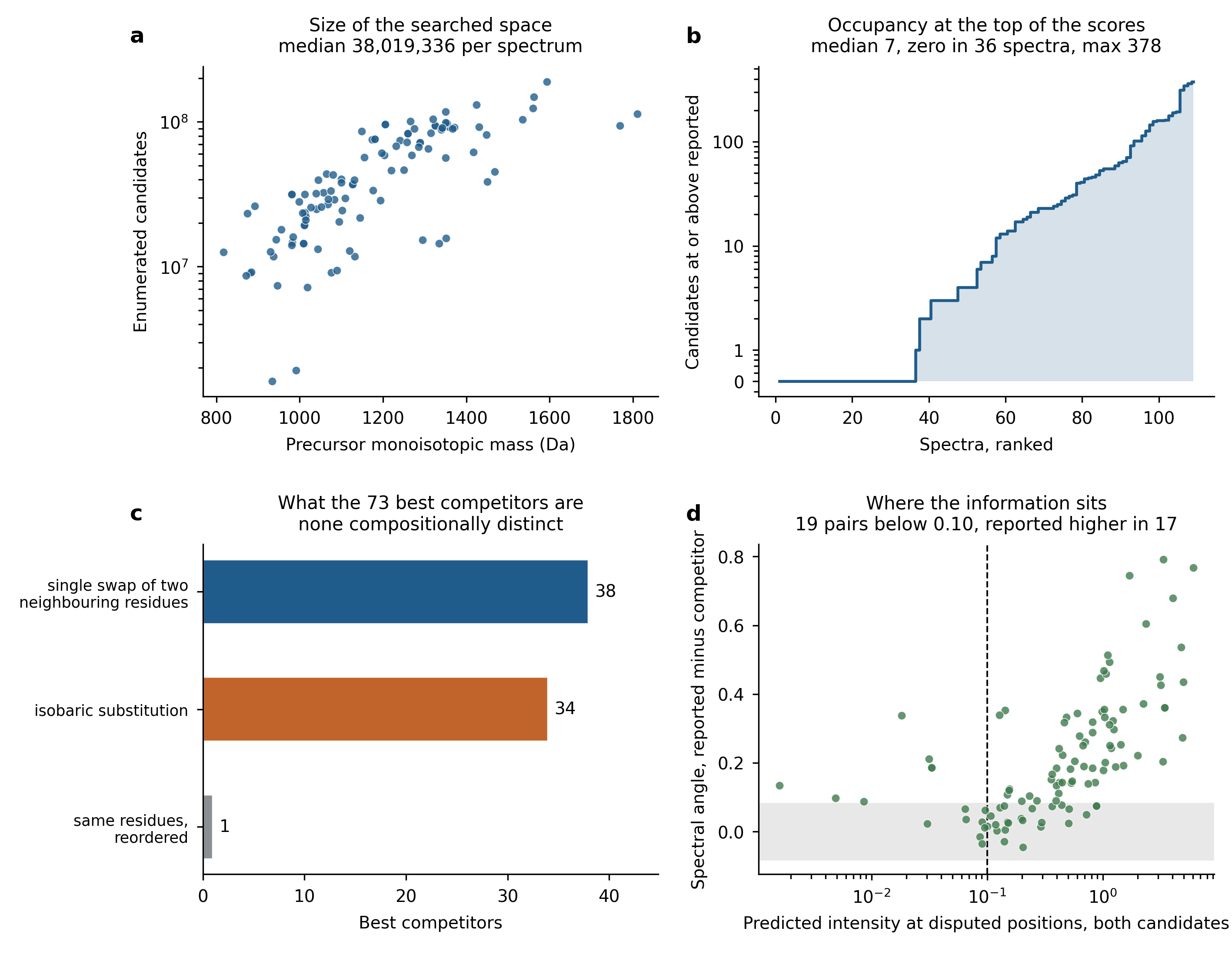

**Supplementary Figure S3.** Structure of the enumerated hybrid-peptide space and of the residual competitors, computed on the original 109 candidate assignments (before the three duplicate scans were removed for the primary verdict; the same set as the PepQuery2 re-evaluation in Supplementary Fig. S4e). (a) Enumerated candidates per spectrum against precursor mass (median 38,019,336). (b) Candidates reaching or exceeding the reported peptide’s matched-ion score (median 7; zero in 36 spectra). (c) Classification of the 73 best competitors after collapsing leucine and isoleucine: 38 single adjacent transpositions, one further reordering, 34 isobaric block substitutions, no compositionally distinct sequence. (d) Summed predicted intensity the two candidates place where they disagree against the resulting spectral angle difference.

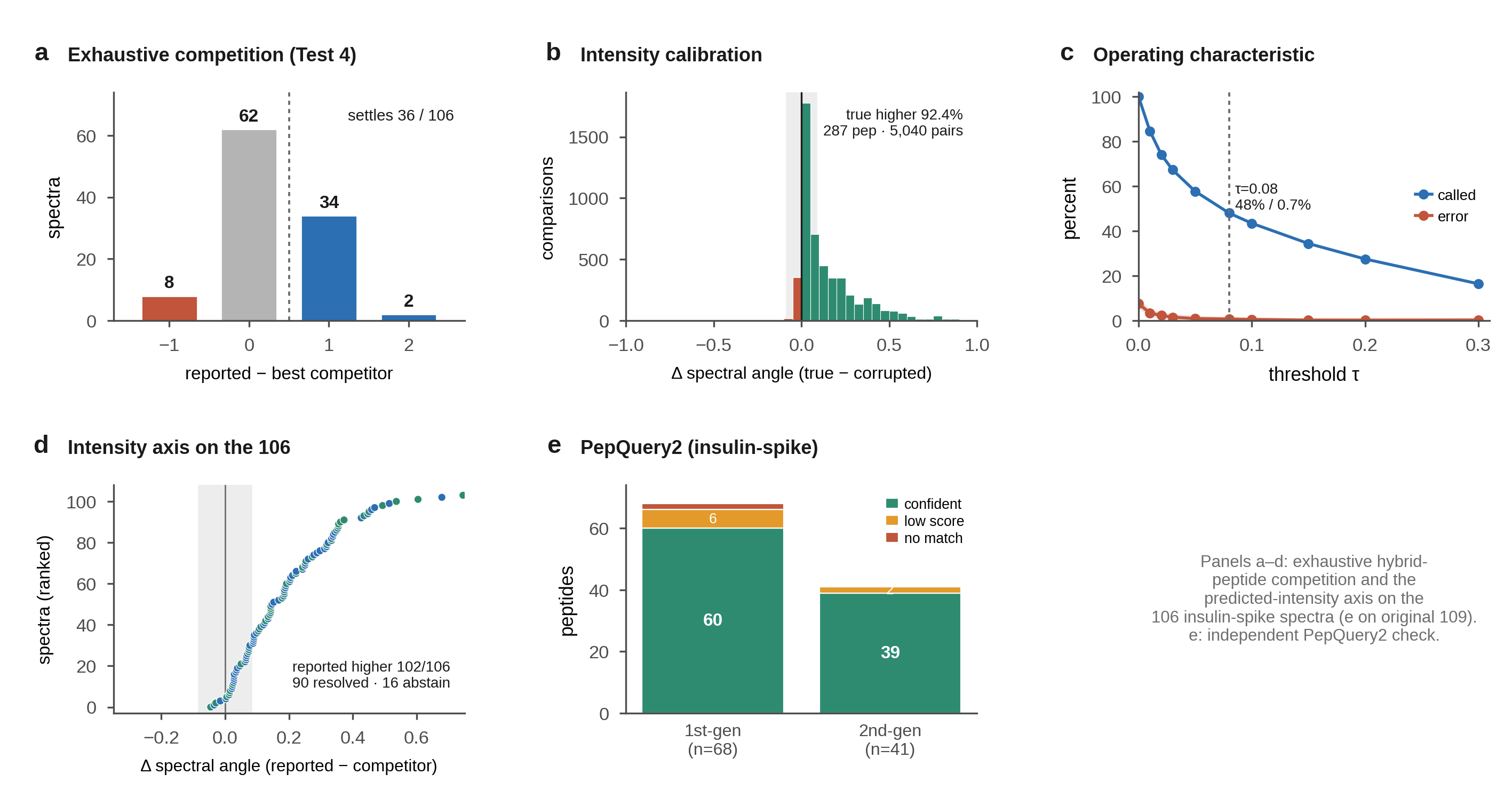

**Supplementary Figure S4. Donor-constrained Test 4 and the predicted-intensity axis on the insulin-spike hybrid peptides.** Secondary evidence for the 106 deduplicated hybrid-peptide spectra; it does not change the primary verdict of 96 retained, 10 unresolved, and 0 rejected. (a) Exhaustive Test 4: the reported peptide minus best donor-hybrid competitor terminal-ion margin over the enumerated candidates (−1 for 8 spectra, 0 for 62, +1 for 34, +2 for 2); the proposed peptide leads on ion count for 36 of 106 and no margin reaches ±3, so Test 4 makes no formal retention or rejection. (b) Predicted-intensity calibration on 287 ProteomeTools peptides of certain truth: the true sequence carries the higher union spectral angle in 92.4 percent of the 5,040 tie-or-lose comparisons; the abstention band |ΔSA| < 0.08 is shaded. (c) Operating characteristic: fraction of comparisons decided and error rate among calls against τ; at the fixed τ = 0.08, 48.0 percent are decided at 0.7 percent error. (d) Predicted-intensity axis on the 106 spectra: the union spectral-angle difference between the reported peptide and its best donor-hybrid competitor, ranked; reported higher in 102 of 106, and among the 70 count-unresolved spectra 54 are resolved for the reported peptide, 16 abstain, and none for a competitor. (e) PepQuery2 independent computational re-evaluation of the original 109 candidate assignments (before the three duplicate scans were removed): first-generation 60 confident / 6 low-score / 2 no-spectrum-match; second-generation 39 confident / 2 low-score.

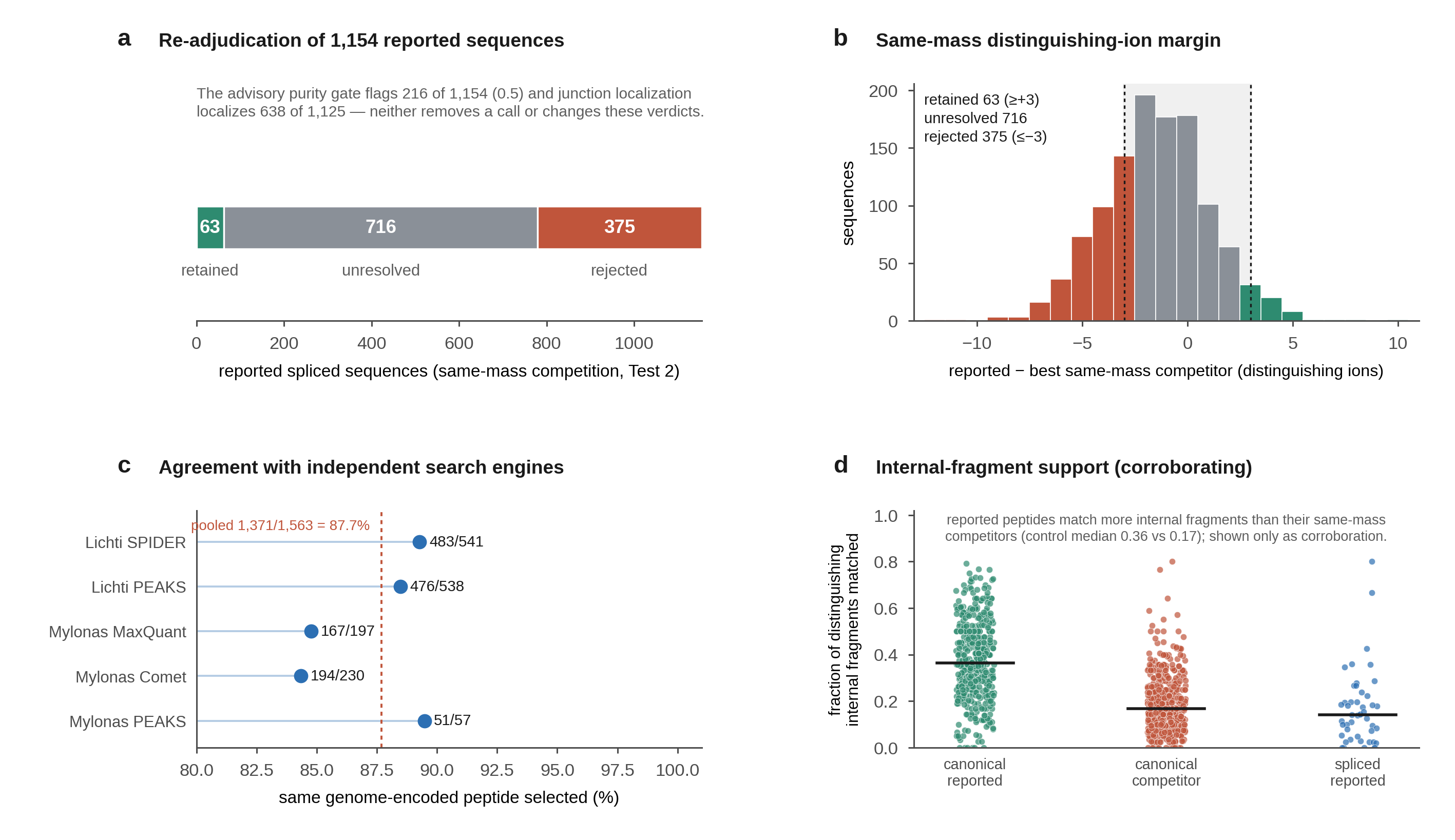

**Supplementary Figure S5. Detail behind the re-adjudication of PXD000394.** (a) Re-adjudication of the 1,154 reported spliced sequences under same-mass competition (Test 2): 53 retained, 718 unresolved, and 383 rejected. The advisory co-isolation purity gate flags 216 of the 1,154 sequences (threshold 0.5) and junction localization localizes 638 of the 1,125 sequences that carry an enumerable junction; neither removes a call or changes these verdicts. (b) Distribution of the same-mass distinguishing-ion margin, defined as the matched distinguishing ions of the reported sequence minus those of its best same-mass genome-encoded competitor; a sequence is retained at a margin of at least +3, rejected at −3 or below, and left unresolved between (shaded band), giving the 53, 718, and 383 of panel a. (c) Agreement with five independent reanalyses (four search engines): for each engine, the fraction of its own reported sequences for which the cascade selects the identical genome-encoded peptide (51 of 57, 194 of 230, 167 of 197, 476 of 538, and 483 of 541; pooled 1,371 of 1,563, 87.7 percent). The engines analyse overlapping spectra and are therefore not mutually independent, so the pooled value is descriptive only. (d) Internal-fragment support, shown only as corroboration outside the primary verdict: reported peptides match a higher fraction of distinguishing internal fragments than their same-mass competitors (control medians 0.36 for the reported peptide against 0.17 for the competitor). No sequence is retained or rejected on this measurement.

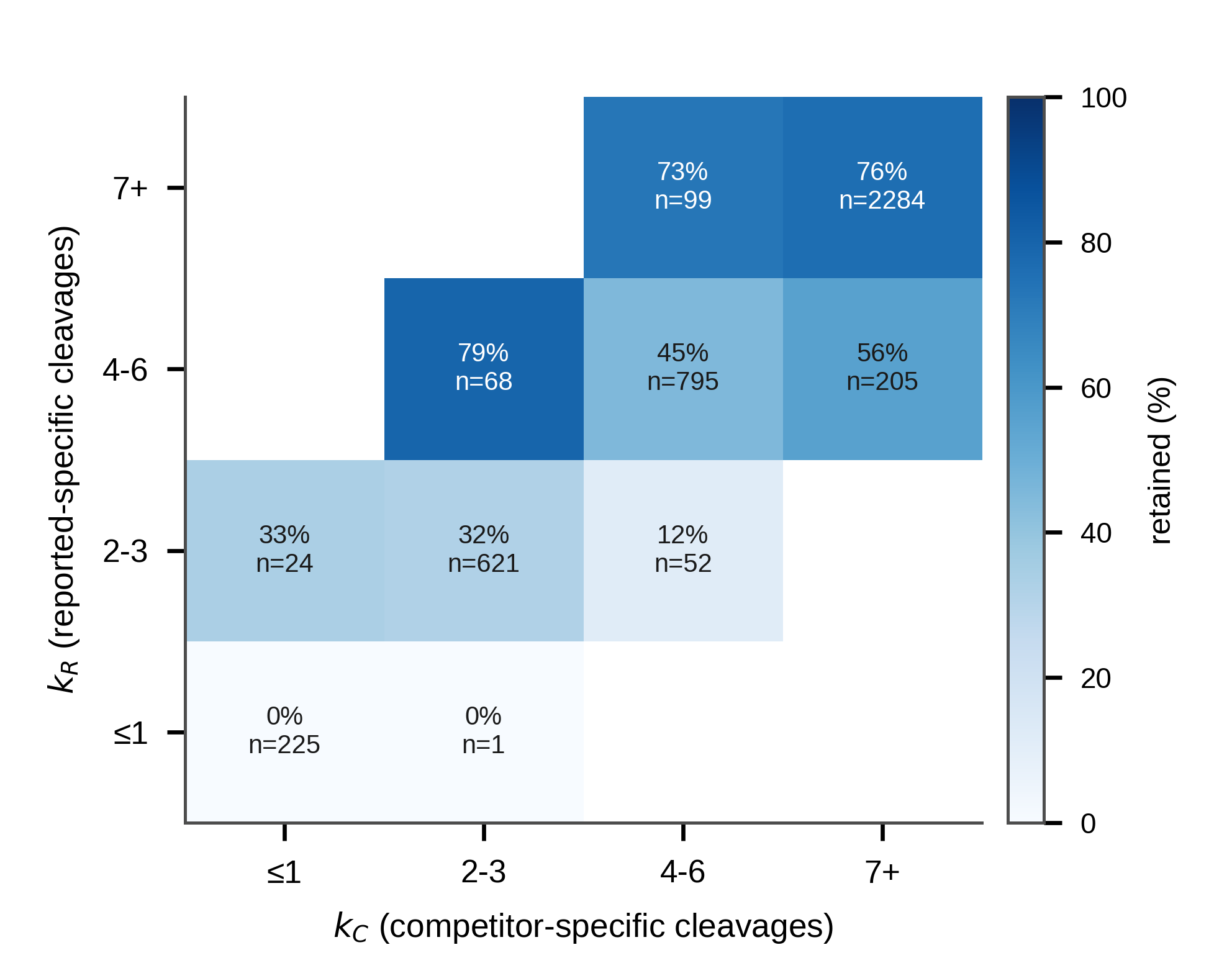

**Supplementary Figure S6. The full (kR, kC) resolving surface on the iBench benchmark.** Retained fraction for all 4,374 spectra binned by kR (internal fragment masses unique to the reported sequence) and kC (unique to the competitor). Retention rises with kR and the competitor’s success rises with kC; the Figure 2d bins are the kK margin of this surface.

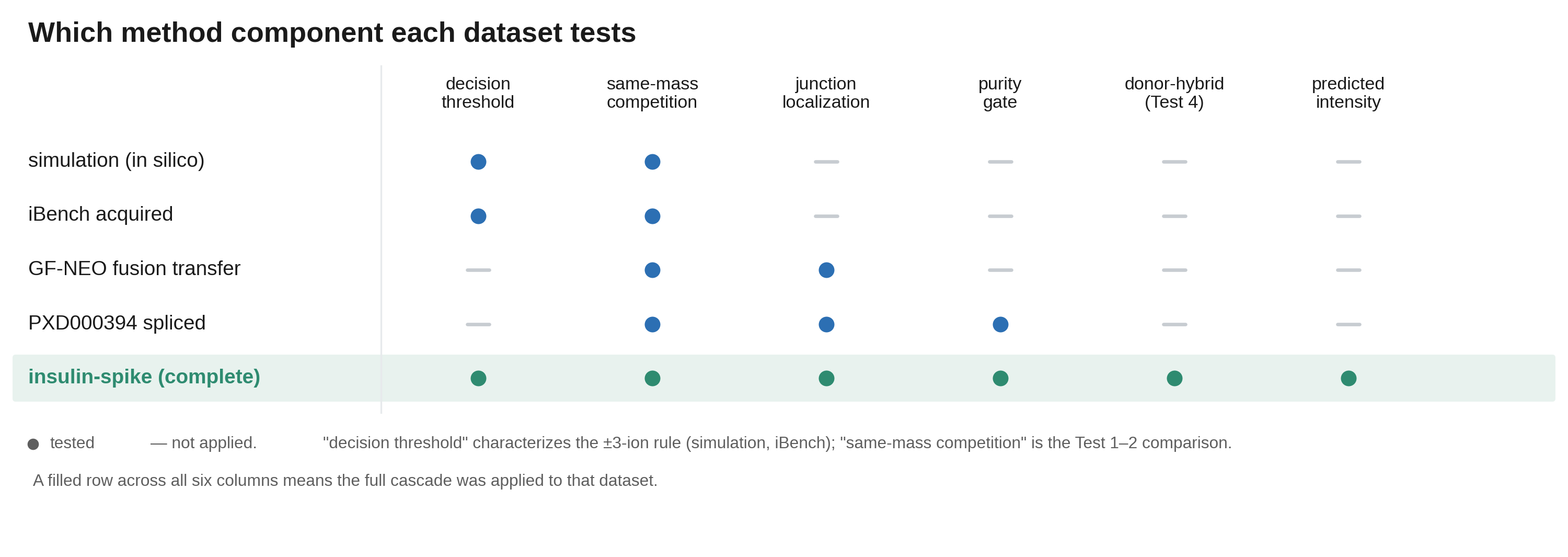

**Supplementary Figure S7. Which method component each dataset tests.** A filled circle marks a component exercised by that dataset and a dash a component not applied. Only the insulin-spike dataset exercises every component: the simulation and iBench benchmark test the decision threshold and the same-mass competition, the GF-NEO transfer adds junction localization, the PXD000394 re-adjudication adds the precursor-purity gate. “Decision threshold” characterizes the three-ion rule where truth is known by construction (simulation, iBench); “same-mass competition” is the Test 1 and Test 2 comparison against proteome-encoded peptides.

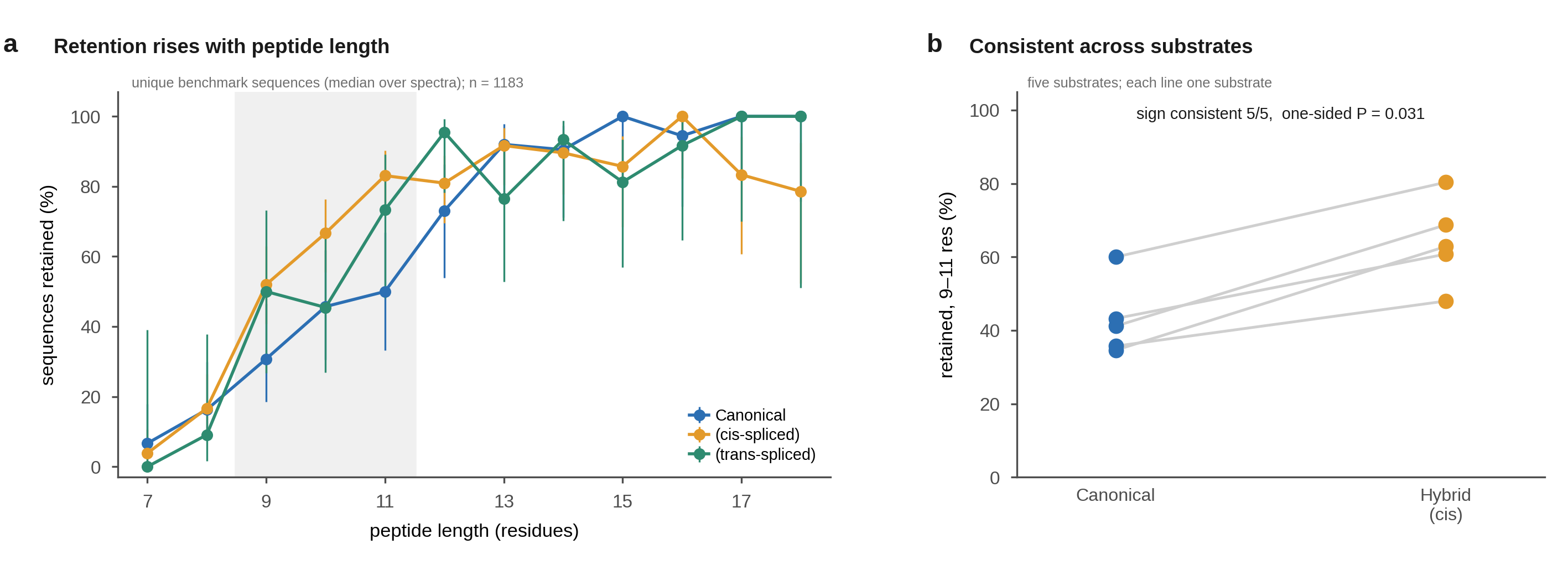

**Supplementary Figure S8. iBench retention by peptide length and by substrate.** Per unique benchmark sequence (median over its spectra, n = 1,183). (a) Fraction retained at a three-ion distinguishing margin against exact peptide length, for canonical, cis-spliced, and trans-spliced peptides; bands are 95% Wilson intervals. (b) Retention at 9 to 11 residues for canonical against cis-spliced peptides in each of the five substrates; the sign is consistent across all five (exact one-sided sign test P = 0.031).

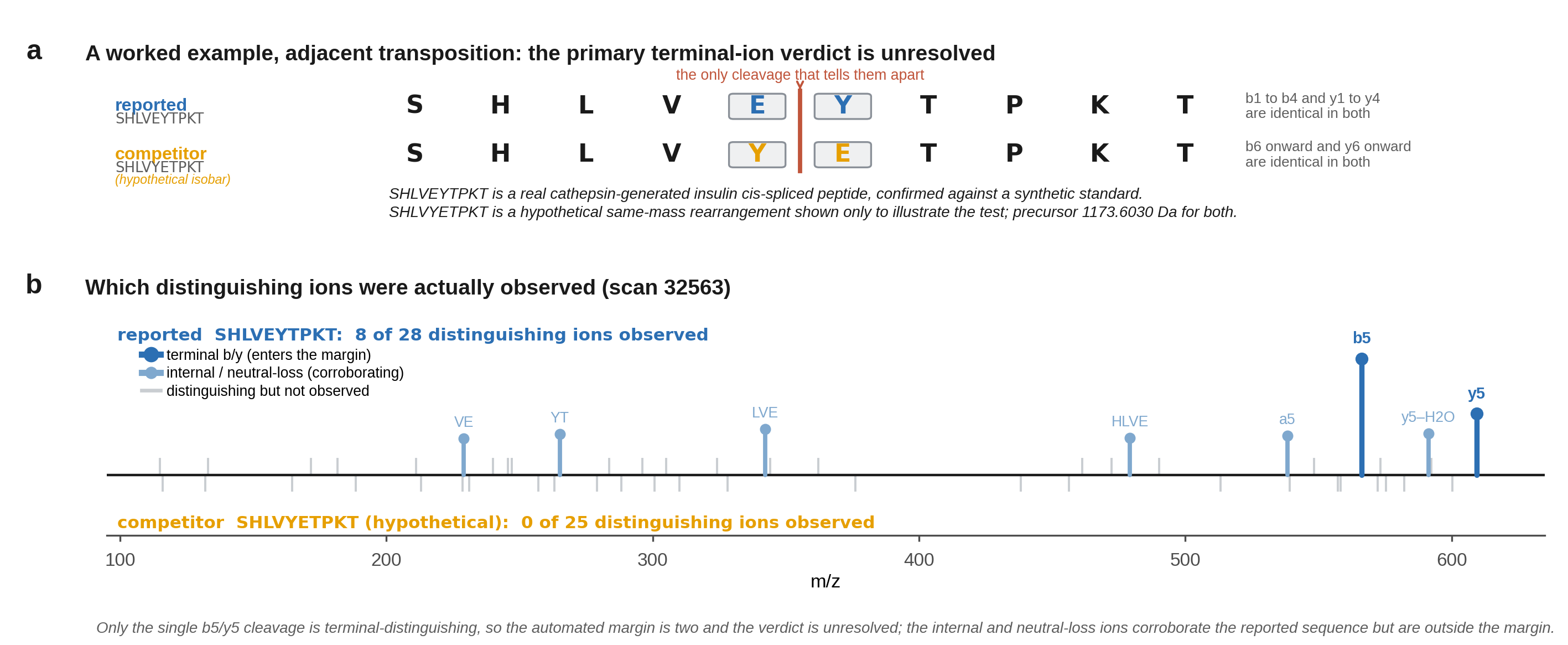

**Supplementary Figure S9. The terminal-ion limit on a single worked example.** SHLVEYTPKT is a real cathepsin-generated insulin cis-spliced peptide, confirmed against a synthetic standard; SHLVYETPKT is a hypothetical same-mass rearrangement shown only to illustrate how the cascade behaves when two isobars differ at a single cleavage. (a) The two sequences as a residue ladder: they are exact isobars (1173.6030 Da) differing by an adjacent glutamate/tyrosine transposition, so a single distinguishing fragmentation separates them and only the terminal b5 and y5 ions distinguish the two. (b) The observed distinguishing ions at scan 32563: eight of the reported peptide’s 28 distinguishing ions were observed and none of the hypothetical competitor’s 25 (Supplementary Tables ST4 and ST5); only the terminal b5/y5 pair enters the three-ion margin, so the automated margin is two and the verdict is unresolved, while the internal and neutral-loss ions corroborate the reported sequence outside the margin.

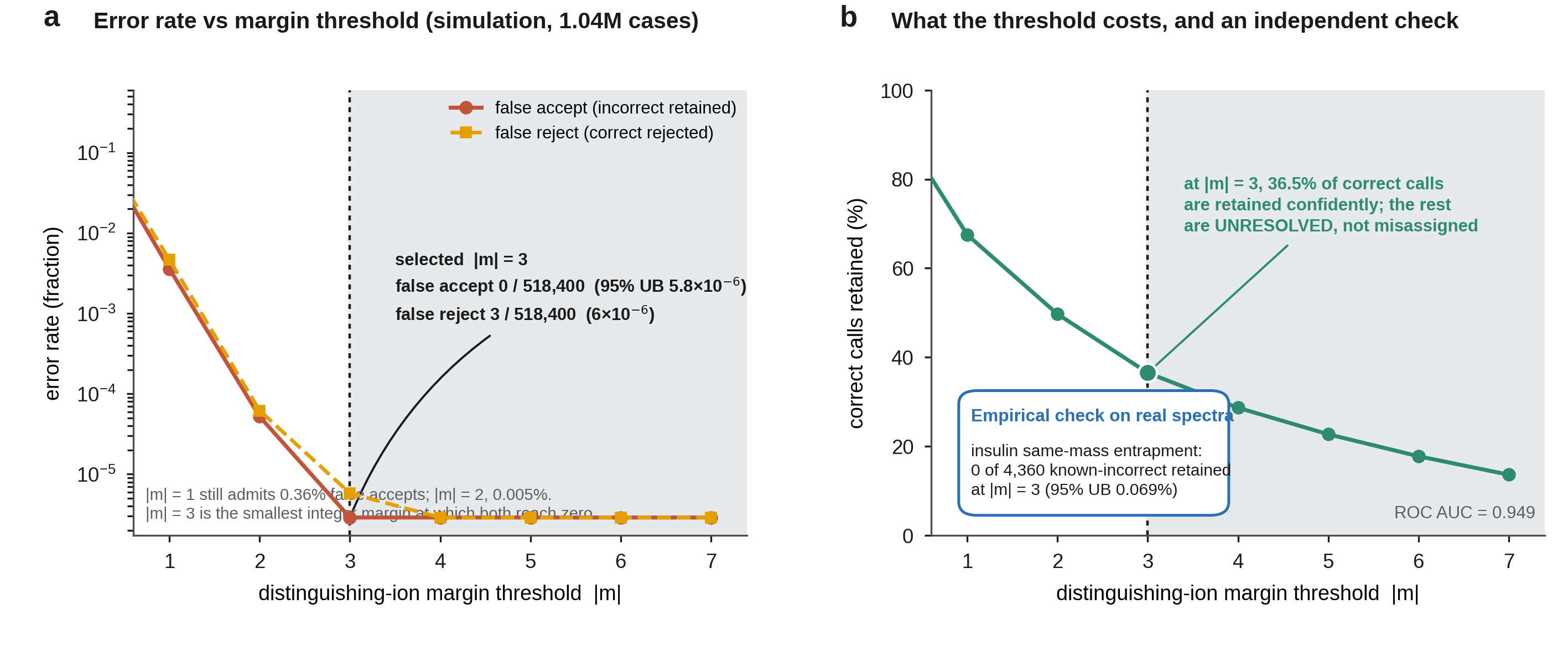

**Supplementary Figure S10. Operating characteristic of the distinguishing-ion margin.** The retention rule (retain at margin m ≥ +3, reject at m ≤ −3) was calibrated on a simulation of 1,036,800 same-mass cases (518,400 known-correct reported peptides and 518,400 known-incorrect same-mass competitors). (a) False-accept rate (a known-incorrect competitor retained) and false-reject rate (a known-correct peptide rejected) as a function of the margin threshold |m|. Both fall monotonically and reach zero at |m| = 3: 0 of 518,400 false accepts (95% upper bound 5.8×10⁻⁶) and 3 of 518,400 false rejects (6×10⁻⁶), whereas |m| = 1 still admits 0.36% false accepts and |m| = 2 admits 0.005%. The threshold of 3 is therefore the smallest integer margin at which both error types vanish, not a chosen heuristic. (b) The cost of the threshold: at |m| = 3, 36.5% of correct calls are retained with confidence and the remainder are classified unresolved rather than misassigned (ROC AUC = 0.949). An independent empirical check on real spectra agrees: in the insulin-spike same-mass entrapment, 0 of 4,360 known-incorrect competitors were retained at |m| = 3 (95% upper bound 0.069%).
